# Identification of anisotropy in chromosome dynamics by principal component analysis using integrated spatial genomics

**DOI:** 10.1101/2024.01.27.577362

**Authors:** Takuya Nara, Haruko Takahashi, Akinori Awazu, Yutaka Kikuchi

**Affiliations:** Graduate School of Integrated Sciences for Life, Hiroshima University, Kagamiyama 1-3-1, Higashi-Hiroshima, Hiroshima, 739-8526 Japan

**Keywords:** Chromosome dynamics, Principal component analysis, Anisotropy, Integrated spatial genomics

## Abstract

Eukaryotic interphase chromosomes maintain a three-dimensional structure within the nucleus and undergo fluctuations. It has been reported that such dynamics are involved in transcription, replication, and DNA repair. However, the analysis of chromosomal dynamics has been limited to high-throughput chromosome conformation capture data, which records the contact frequencies between chromosomal regions and lack direct information about the dynamic. Herein, we investigated chromosome fluctuations as polymers based on experimental data from sequential fluorescence *in situ* hybridization (seqFISH)+ using a multiomics methodology. To describe the principal modes of chromosome fluctuations, we applied principal component analysis to the three-dimensional structure information of single chromosomes in 446 mouse embryonic stem cells (mESCs) obtained from seqFISH+ data analysis for spatial genomics and signals of nuclear factors (SNFs: histone marks, repeat DNAs, and nuclear compartments). We found that chromosome fluctuations exhibit both isotropic and anisotropic modes. The properties of anisotropy in chromosome fluctuation vary among chromosomes and appear to depend on the interaction between repeat DNAs on the chromosomes and nuclear compartments. Furthermore, our principal component analysis revealed anisotropic chromosome fluctuations before and after the mitotic phase, specifically when chromosomes adopt a spindle-like shape. This result suggests the potential involvement of anisotropic chromosomal fluctuations in the transition of nuclear organization during the cell cycle. Our results represent the first study to elucidate the dynamics of chromosomes as polymers based on real multiomics data.

## Introduction

Although the eukaryotic nucleus is a subcellular organelle surrounded by a lipid bilayer membrane, it does not contain organelles with membrane (1-4). Recent advances in genomics and optical observation methods have revealed that the interphase nucleus is compartmentalized by membrane-less organelles called “biomolecular condensates,” where the specific protein and nucleic acid, primarily RNA, molecules are concentrated (1-4). The subnuclear bodies, such as the nucleolar, speckles, and Cajal bodies, are known as types of biomolecular condensates in the interphase nucleus (2-4). In addition to biomolecular condensates, each chromosome forms a highly compartmentalized architecture, named “chromosome territory” (5, 6). Recent super-resolution imaging has revealed that chromatin fibers extending outside the chromosomal territory interact with various biomolecular condensates located in the interchromosomal space and nuclear lamina/pore complexes at the nuclear periphery (6-8). Such interactions between chromatin fibers and biomolecular condensates have been proposed to be correlated with transcription, RNA splicing, and DNA replication (6-8).

In the interphase, chromatin is hierarchically self-organized to form three-dimensional chromatin fiber-like chromatin loop domains/compartments and topologically associated domains, and these fibers exhibit irregular and dynamic liquid-state properties (9, 10). Chromosome conformation capture (3C)-based, fluorescence *in situ* hybridization (FISH) technologies and the methodologies based on the bacteriophage MS2/PP7 coat-binding protein or tagged dead CRISPR associated protein 9 have been used to understand the chromatin organization and dynamics (11, 12). The three-dimensional (3D) folding and fluctuations in chromatin are involved in the regulation of DNA-related processes, such as transcription, replication, and DNA repair, by linking chromatin fibers to interactions with biomolecular condensates and the nuclear periphery (6, 13). However, the folding and fluctuation of chromatin are highly complex processes, making the interpretation of chromatin/chromosome dynamics from a combination of 3C data, especially high-throughput chromosome conformation capture (Hi-C) and imaging results, challenging. Therefore, computer simulations have also been used by applying polymer physics, considering chromosomes as high-molecular-weight polymers (14, 15). The increase in data volume and improvement in resolution through Hi-C and imaging technologies have greatly advanced research on quantitative computational models for predicting chromatin folding and chromosome dynamics (10). However, to data, few computational studies have been conducted to analyze chromatin/chromosome dynamics based on experimental data, such as FISH imaging and Hi-C without simulations.

Unlike studies on chromatin folding and dynamics, the higher-order structure and dynamics of proteins have mainly been studied using quantitative techniques such as X-ray crystallography and nuclear magnetic resonance (NMR), which are often unable to provide a complete trajectory of protein dynamics (16, 17). Structural bioinformatics methods, such as molecular dynamics (MD) simulations complement the limitations of experimental techniques, allowing for the prediction of protein dynamics at the atomic level (16, 17). In addition to MD, normal mode analysis (NMA) and principal component analysis (PCA) have been used to determine large-scale amplitude motions in proteins (18, 19). The NMA/PCA method provides information on protein fluctuations using multiple distance matrices related to molecular shapes obtained using techniques such as MD and NMR (18, 20, 21). The eigenvalues and eigenvectors obtained through the PCA represent the amplitude and direction of the fluctuations corresponding to the modes of the principal components (19, 22). The application of PCA to protein fluctuation analysis revealed that the direction of low-frequency protein fluctuation is not isotropic but rather anisotropic, with each functional domain of the proteins fluctuating in a coherent direction (18, 19). PCA and many related methods can be widely applied to the analysis of protein dynamics and the analysis of biomacromolecule dynamics, including chromosomes, as long as the position data of biomacromolecule components are obtained (18, 19).

With the improvements in single-cell sequencing technology and FISH methods, many multiomics (such as the genome, epigenome, and transcriptome) analysis techniques with spatial information at the single-cell or nuclear level have recently been reported (23, 24). More recently, FISH imaging and multiomics methods have been developed for identifying the location of DNA loci, transcripts, nuclear structures/compartments, including biomolecular condensates and nuclear membranes, and histone modifications within a single nucleus (23, 24). DNA seqFISH+, a multiomics method, provides 3,660 chromosome loci, along with 17 marks of chromatin and nuclear structures/compartments and 70 RNAs in single nuclei of mouse embryonic stem cells (mESCs) (25). Using the data obtained through the DNA seqFISH+ method allows the analysis of the spatial relationships between positions of chromosome regions and signals of nuclear factors (SNFs: histone marks, repeat DNAs, and nuclear compartments) within a single nucleus of mESCs. In this study, we applied PCA, similar to mode analysis of protein dynamics, to elucidate chromosome dynamics based on experimental data, specifically DNA seqFISH+ data.

## Results

### Comparison of chromosome fluctuations analyzed using PCA and the spring-bead model

We attempted to describe chromosomal fluctuations in the nuclei of mESCs using experimental seqFISH+ data as a spatial genomics data set. For this purpose, we extracted a set of fluorescent probe dots derived from a single chromosome in each nucleus (Fig. S1, see *Materials and Methods*) and obtained sets of distance matrices of fluorescence probes for each type of chromosome (chromosomes 1-19 and X) based on the 3D coordinates of the extracted fluorescence probes from single chromosomes (see *Materials and Methods*). The eigenvalues and eigenvectors were calculated using PCA for the set of distance matrices, and the index of fluctuation amplitude (IFA) was examined for each chromatin bin for each type of chromosome (see *Materials and Methods*). We found a significant and positive correlation between the IFA and the genomic distance from the chromosome center for most chromosomes, suggesting a high degree of freedom in the fluctuation at chromosome ends (Fig. S2).

Subsequently, we compared the IFA in Fig. S2A with the results of the chromosome fluctuation simulations based on the contact matrices within the chromosomes derived from the Hi-C data. We selected the PHi-C2 method, which was developed to bridge the gap between live-cell imaging and the Hi-C method, as the simulation method for comparison (26, 27). The PHi-C method formulates that the contact matrices stored in the Hi-C data represent a function of variance concerning the distance distribution between chromosomal domains, and it is designed for dynamic simulations of chromosomes (26, 27). In analyzing chromosomal fluctuations using the PHi-C2 method, a complex compliance for each binned chromosomal region was obtained, representing the ease of deformation when an external force is applied to the spring-bead model of the chromosome. We performed a comparative analysis of the complex compliance obtained using the PHi-C2 method and IFA. In this comparison, it is impossible to align the PHi-C2 and seqFISH+ data using PCA based on a unified scale for the radial frequency variable ω, which is crucial for studying fluctuations. Therefore, for PHi-C2, we investigated three conditions of ω in PHi-C2: 10^-1^, 10^-2^, and 10^-3^, while assuming a constant radial frequency (ω) for the chromosomal fluctuations of seqFISH+ data studied using PCA. As the value of χο, the radial frequency variable of PHi-C2 (28), decreases, meaning that the timescale for investigating fluctuations becomes larger, the Pearson correlation coefficient (PCC) between the IFA and complex compliance obtained using the PHi-C2 method increases (Fig. 1). However, even for ω = 10^-3^, which correlated best with the IFA among the radial frequency variables of the PHi-C2 method examined, the median PCC among chromosomes was approximately 0.1 (Fig. 1). These results suggest that dynamic simulations of chromosomes are insufficient when only the variance concerning the distance distribution between intrachromosomal domains is considered. This suggests the importance of considering the following two aspects: (1) the examination of interactions between chromosome regions and nuclear compartments and (2) the examination of interactions between chromosomes.

**Figure 1.**
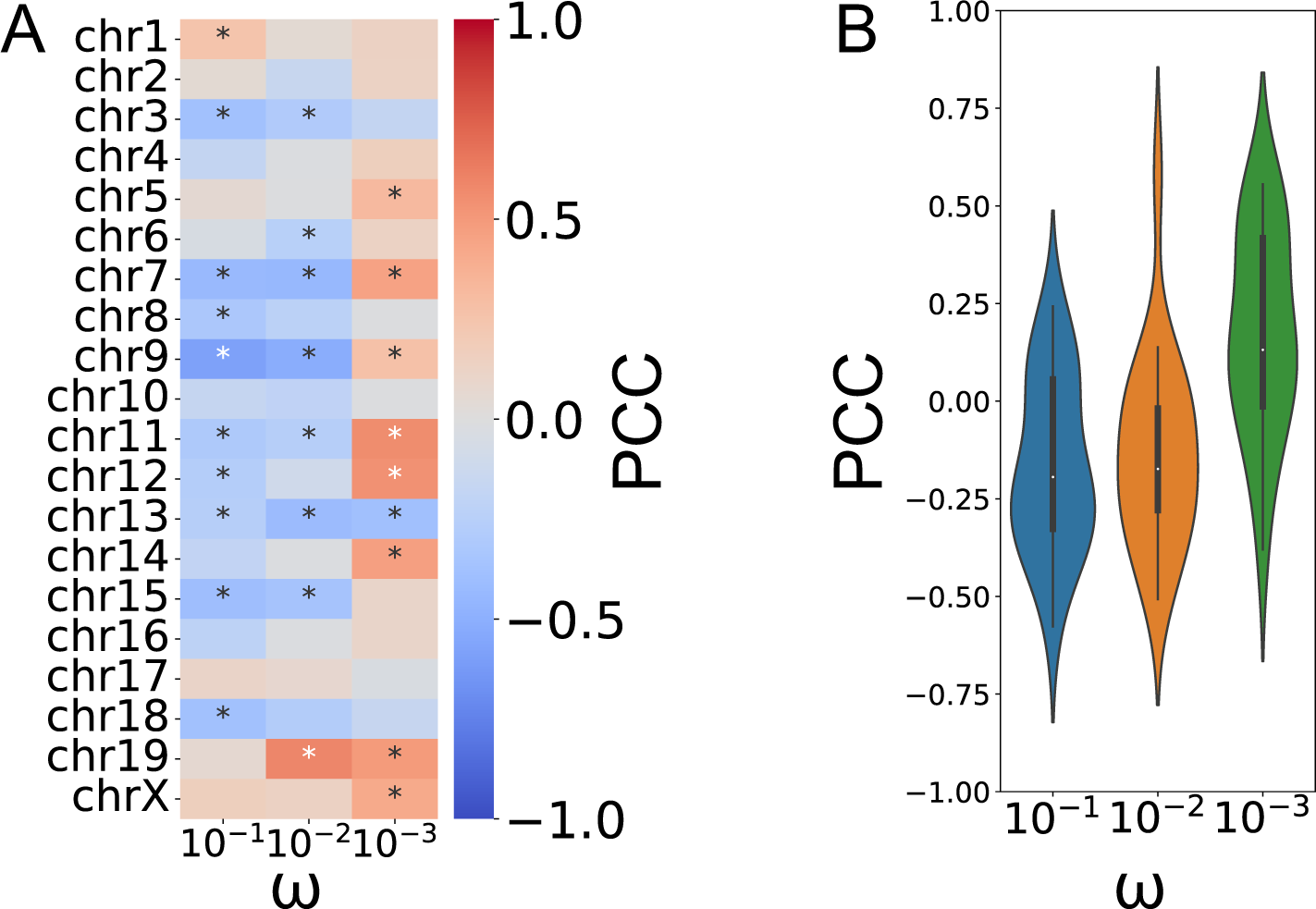
Comparison between the fluctuation amplitude and the complex compliance simulated by PHi-C2 along chromosomes. (A) A heatmap shows the Pearson correlation coefficient (PCC) between the index of fluctuation amplitude (IFA) and the complex compliance for each chromosome and each radial frequency parameter, ω. The following is the number of bins: chromosome 1 (A: N = 178), 2 (B: N = 185), 3 (C: N = 152), 4 (D: N = 149), 5 (E: N =140), 6 (F: N = 154), 7 (G: N = 142), 8 (H: N = 113), 9 (I: N = 134), 10 (J: N = 130), 11 (K: N = 125), 12 (L: N = 107), 13 (M: N = 119), 14 (N: N = 100), 15 (O: N = 100), 16 (P: N = 91), 17 (Q: N = 83), 18 (R: N = 82), 19 (S: N = 65), and X (T: N = 120). \**P* < 0.01 using the Pearson test. (B) A violin plot shows the distribution of PCC between IFA and the complex compliance for each chromosome, and each radial frequency parameter, ω (N = 20 types of chromosomes).

### Correlation analysis of IFAs and SNFs based on experimental data

To explore the relationship between IFAs and SNFs, we used seqFISH+ of mESCs as an integrated multiomics data set. SNFs derived from the fluorescence signals analyzed using seqFISH+ are listed in Table S1. IFA was positively correlated with the frequency of contact with telomeres, as well as with the constitutive heterochromatin markers H3K9me3, H4K20me3, and mH2A1, which are found in the telomeric region (Fig. 2A). We next examined the relationship between IFAs of chromatin bins and the proximity to nuclear compartments (Fig. S3 and Table S2), by defining the location of nuclear compartments based on the distribution of SNFs in the seqFISH+ data (see *Materials and Methods*). A Gaussian mixture model (GMM) was employed because the Bayesian information criterion (BIC) allows the number of clusters to be considered most appropriate for the data. We adopted the number of clusters set for the definition of nuclear compartments with GMM as 60, which corresponded to the minimum BIC when considering cluster numbers ranging from 2 to 60 (Fig. S3A). Sixty clusters were used to define the nuclear compartment from the seqFISH+ data, and the 60 clusters obtained using GMM were interpreted using signal values corresponding to each SNF (Table S2). We found an almost negative correlation between the IFA of chromatin bins and “proximity to nuclear compartments” (Fig. 2B), suggesting that presence of nuclear compartments suppresses the amplitude of chromosome fluctuations.

**Figure 2.**
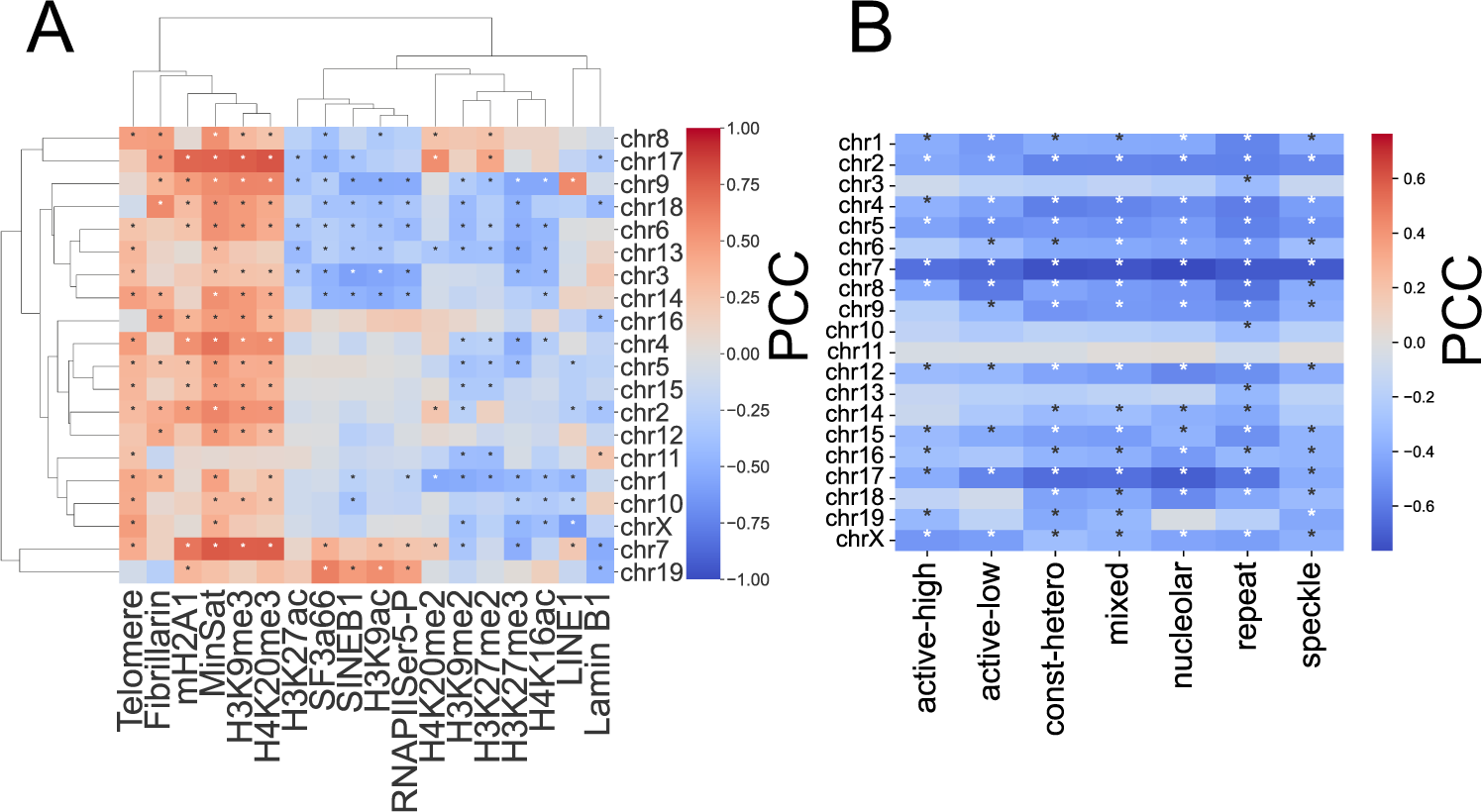
The nature of the distribution of IFA along chromosomes. (A) A heatmap shows the PCC between IFA and the frequency of contact between chromosome bins and signals of nuclear factors (SNFs) (N = 446 cells). SNFs derived from the seqFISH+ were considered (see Tab. S1 and Materials and Methods). \**P* < 0.01 using the Pearson test. (B) A heatmap shows the PCC between IFA and the minimum distance from chromatin bins to nuclear compartments (N = 446 cells). Nuclear compartments based on the fluorescent signals (See Materials and Methods, Fig. S2, and Tab. S2) were considered. \**P* < 0.01 using the Pearson test.

### The distribution of eigenvectors, which represents the direction of the chromosome fluctuations

In addition to IFAs, we further investigated whether there were variations in the direction of chromosomal fluctuations among the PC modes. To visualize the direction of chromosomal fluctuations, we embedded the first three eigenvectors obtained using PCA into the average 3D structure of the chromosomes (see *Materials and Methods*). Our results showed that the first principal component (PCmode1) fluctuated in the direction corresponding to the contraction and relaxation of the chromosome (Fig. S4). However, eigenvectors were observed in the second and third principal components (PCmode2 and PCmode3), which showed fluctuation directions such that when most parts of the chromosomes contracted, some chromosomal regions were pushed in the opposite directions (Figs. 3A and S4). We then examined whether there were differences in the distribution of the eigenvectors based on the orientation of the chromosome (from the p-arm to the q-arm). In other words, we investigated whether anisotropy existed in PCmode1, PCmode2, and PCmode3 based on the orientation of the chromosome. In this analysis, we used the PCC between the matrix of the eigenvector (“original-matrix”) with the pairwise coordinates of chromosome bins and the matrix (“flipped-matrix”) obtained by flipping the “original-matrix” vertically and horizontally (Fig. 3B). In this case, the PCC approaches 1 if the eigenvector is isotropically distributed based on the chromosome orientation and approaches -1 if it is anisotropically distributed. As shown in Fig. 3C, the distribution of PCmode1 was isotropic, whereas the distributions of PCmode2 and PCmode3 exhibited weaker isotropic properties than PCmode1.

**Figure 3.**
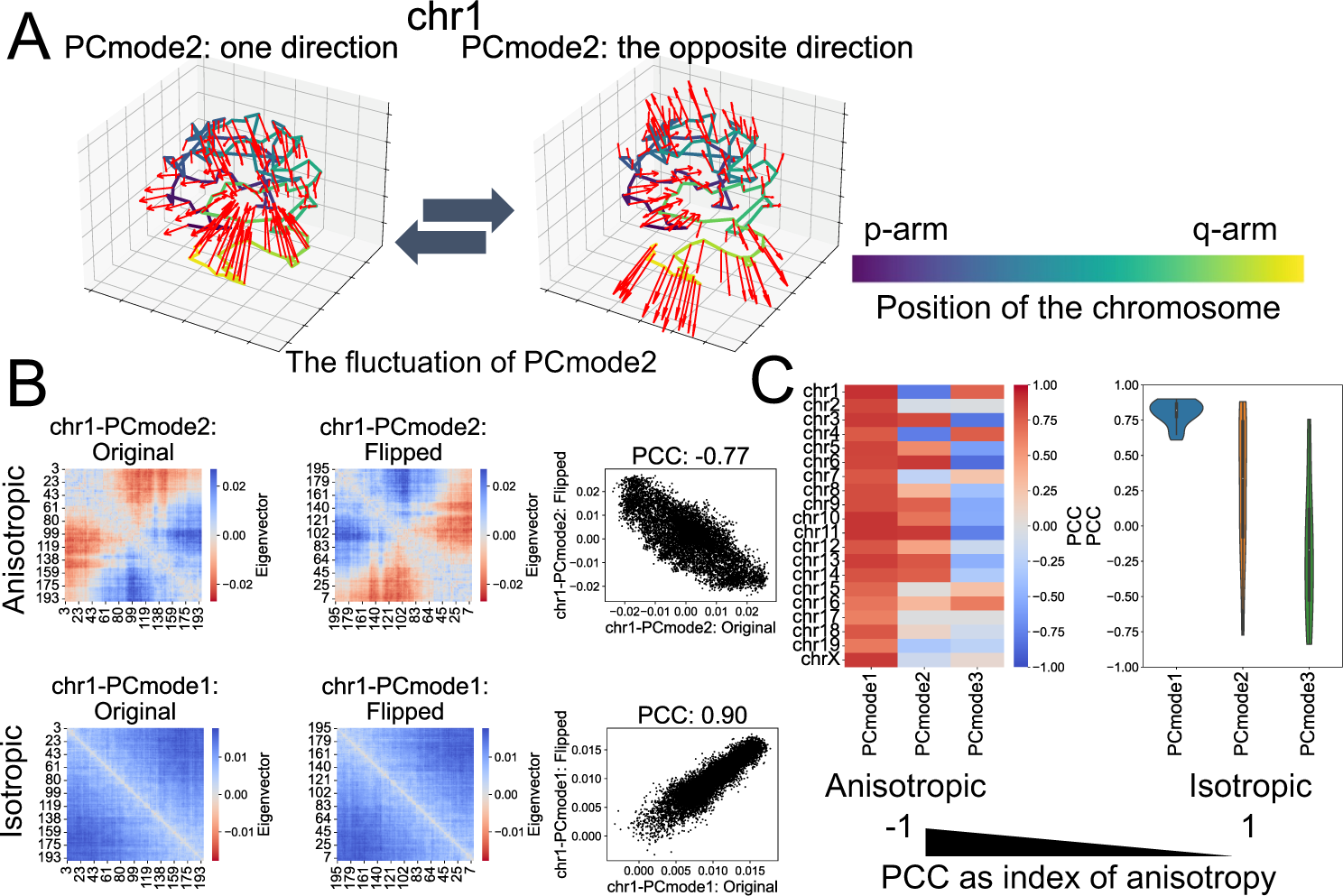
Quantification of anisotropy in the distribution of eigenvector. (A) Enlarged images show the PCmode2 of chromosome 1 (chr1) fluctuation. The left and right panels correspond to one and the other direction of fluctuation. The color bar shows the relative position of bins along the chromosome (i.e., The darkness of the color scale means the proximity from the end of the p-arm). (B) The example of quantifying anisotropy in chromosome fluctuation using correlation analysis. 1st step: Construct the eigenvector matrix corresponding to the coordination of the pairwise distance matrix of the chromosomes (left panels). 2nd step: Convert the eigenvector matrix by flipping upside down and leftside right (middle panels). 3rd step: Find PCC as the index of anisotropy in the eigenvector of chromosome fluctuation (right panels). X- and Y-axes indicate the genomic coordination in the chromosome (Mb: Left and right panels). (C) The PCC is the index of anisotropy in the eigenvector of chromosome fluctuation (N = 20 types of chromosomes).

### The relationship between fluctuation and shape of a single chromosome for each cell

To reveal when anisotropic fluctuations, as identified based on eigenvectors, occur in chromosome organization, we analyzed the relationship between the chromosome structure and the PC scores of each PC mode. The seqFISH+ data used in this study were obtained from mESCs whose cell cycles were not artificially controlled (25). Therefore, it was assumed that the cells sampled in the data were randomly selected from various processes of the entire cell cycle. In addition, the eigenvectors of PCmode1 in the chromosome fluctuations were isotropic (Fig. 3C), suggesting that PCmode1 is associated with fluctuations involved in the condensation/decondensation of chromosomes. We first examined whether anisotropic chromosome fluctuations occurred when the chromosomes condensated or decondensated by focusing on the probability distribution of the PC scores of PCmode1 and PCmode2/PCmode3 for each chromosome (Fig. S5). In this analysis, we divided the PC scores of PCmode2/PCmode3 into two groups based on the sign of the PC scores of PCmode1 and examined the ratio of variances between the two groups. The ratio of variances (’log2R on variance’) was more than twofold for all types of chromosomes (Fig. S5), indicating that anisotropic fluctuations represented by PCmode2/PCmode3 are observed when chromosomes are in either a condensed or decondensed state. Subsequently, to investigate how PCmode1 of each chromosome corresponded to the condensation/decondensation of chromosomes, we examined the correlation between the PC scores of PCmode1 and the variance of distance matrices of single chromosome (VDMSC). We found a high correlation (|PCC| > 0.8) between PCmode1 and VDMSC for all types of chromosomes and observed that in the group with larger variances in the PC scores of PCmode2/PCmode3, VDMSC tended to be larger, regardless of the chromosome type (Fig. S5). Furthermore, when examining how VDMSC corresponds to the condensation or decondensation of chromosomes, we found that when the VDMSC of each chromosome was large, the chromosomes tended to have a spindle-like shape, whereas they exhibited a spherical shape when the VDMSC was small (Figs. 4A, 4B, and S6). These results suggest that anisotropic fluctuations, represented by PCmode2/PCmode3, are more likely to be observed when chromosomes resemble condensation rather than decondensation.

**Figure 4.**
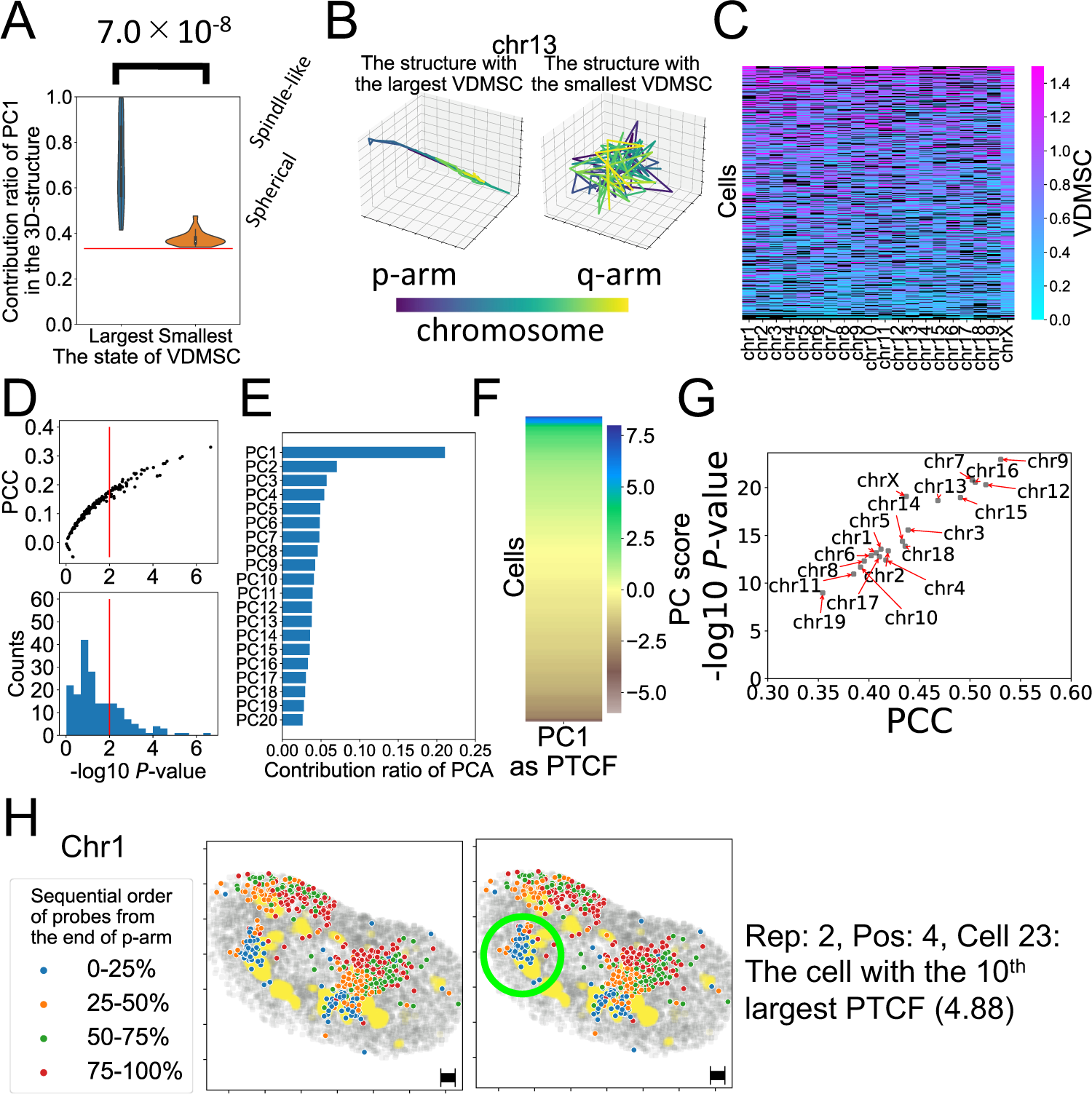
The representative states of chromosomes and cells correspond to the VDMSC. (A) The contribution ratio of PC1 for the structures of a single chromosome with the largest and smallest VDMSC (N = 20 structures). The value on the bracket means the result of the Student’s *t*-test as a *P*-value. (B) The structures of single chromosomes with the largest and smallest VDMSC. A color bar shows the relative position along chromosomes (i.e. The darkness of color scale means the proximity from the end of p-arm). (C) The mean of VDMSC from each cell (N = 427 cells). Black panels indicate the missing values. (D) Correlation of VDMSC between each type of chromosome. Red lines indicate the *P*-value of the Pearson test = 0.01. (E) The contribution ratio of PCA on the matrix of VDMSC from each cell in Fig. 4C. (F) The PTCF that was calculated as the first PC score of PCA on the matrix of VDMSC in Fig. 4C serves as the index of chromosome condensation/decondensation in each cell. (G) The correlation between PTCF and the mean of VDMSC for each cell. (H) The structures of Chr1 in the cell with the 10th largest PTCF. Blue, orange, green, and red dots correspond to genomic coordination of the probes from 0-25%, 25-50%, 50-75%, and 75-100% from the end of p-arm. Yellow dots indicate the position of fibrillarin, a nucleolar marker. Green circles in the left panel indicate the chromosome regions that correspond to the anisotropic parts of chromosome structures. Black bars indicate 1µm for X- and Y-axes in seqFISH+ data.

The spindle-like or spherical shapes of the chromosomes obtained from our analysis were presumed to reflect their cell cycle-dependent condensation/decondensation shapes. Therefore, VDMSC between chromosomes was expected to be synchronized for each cell. To test this hypothesis, we examined the correlation of the average number of VDMSC between chromosomes in each cell. Our results showed a positive correlation between VDMSC for all chromosome types (Figs. 4C and 4D), suggesting that the VDMSC of each chromosome type synchronizes across cells. Furthermore, to investigate the synchronization of VDMSC among cells, we defined an index corresponding to the condensation/decondensation state of chromosomes for each cell, termed the pseudo-trajectory of chromosome fluctuation (PTCF). The PTCF was generated as the first PC score obtained by applying PCA to the matrix of the averaged VDMSC of each chromosome type in each cell, as shown in Fig. 4C (Figs. 4E and 4F). Consistent with this definition, there was a positive and significant correlation between PTCF and the average VDMSC of each chromosome type in each cell (Fig. 4G). Some single-chromosome structures in the cells with higher PTCF values exhibited anisotropy (Figs. 4H and S7). This suggests that chromosomes exhibit anisotropic fluctuations before and/or after the mitotic phase (M phase) when they condense into spindle-like shapes. Furthermore, by applying PCA to the 3D distribution of probes derived from single chromosomes to examine the divergence of single-chromosome shapes, we found that some single chromosomes adapted to a bent anisotropic structure when in a state closer to a spindle-like morphology (Figs. S8A and S8B). In addition, cells with higher PTCF values tended to have spindle-like shapes (Fig. S8C).

### Exploration of nuclear factors (histone marks, repeat DNAs, and nuclear compartments) related to the direction of chromosome fluctuations

To elucidate the characteristics of the chromosomal regions constituting anisotropic fluctuations in PCmode2/PCmode3, we focused on the signs of the eigenvectors for each PC mode associated with the chromosomal bins (Fig. 5A and *Materials and Methods*). The investigation of chromosome fluctuations using PCA in this study was based on the distances between probes obtained from seqFISH+ (Figs. 5A① and 5A②). Therefore, the eigenvectors of each PC mode describe the direction of the principal variances in distances between probes related to chromosome fluctuations (Figs. 5A③ and 5A④). In other words, the signs within the eigenvectors of each PC mode reflect the direction of fluctuation, i.e., the pattern of increase or decrease in distances between chromosome regions in that PC mode (Fig. 5A④). We defined the direction corresponding to the less frequent sign when comparing the number of positive and negative values within each eigenvector as the minor direction (MND) (Fig. 5A④). For the eigenvectors of PCmode1 in all chromosomes, the ratio of MND was highly unbalanced (ratio < 1%) (Figs. S9 and S10), suggesting that the direction of chromosomal fluctuation in the PCmode1 was isotropic within the eigenvector, that is, a cooperative increase or decrease in the pattern of distances between intrachromosomal regions. For the eigenvectors of the PCmode2 and PCmode3, the ratio of MND was balanced (ratio: 30-50%) compared with that of PCmode1 (Figs. S9 and S10), suggesting that the direction of chromosome fluctuation in PCmode2 and PCmode3 is anisotropic within the eigenvector, that is, a non-cooperative increase or decrease pattern of distances between intrachromosomal regions.

**Figure 5.**
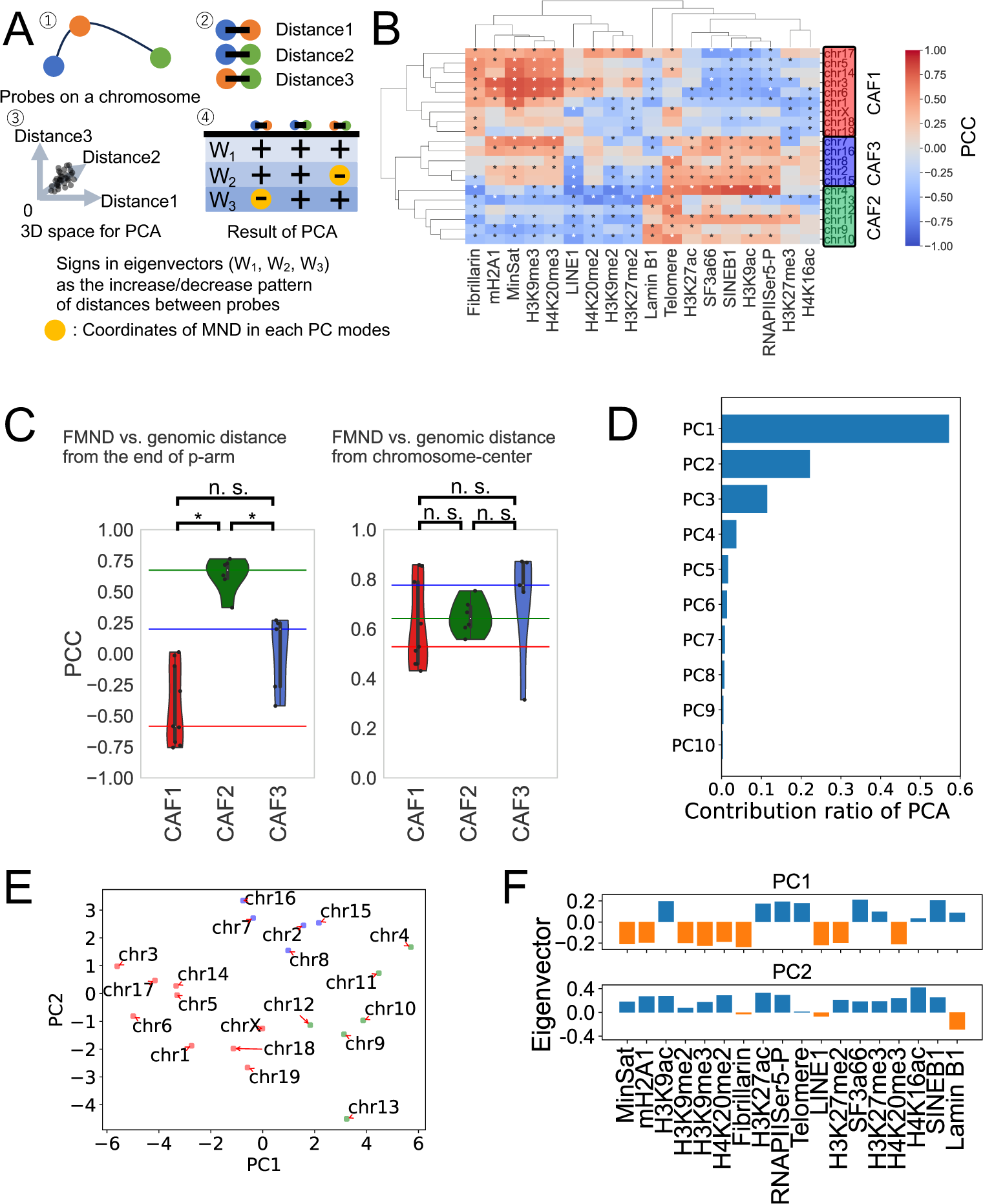
Comparison between FMND along chromosomes and the distribution of SNFs in nuclei. (A) Explanation of the meaning of the signs of eigenvectors. Based on the position of probes on a chromosome (①), we measured the distances between probes (②). By applying the PCA on the multidimensional space corresponding to the number of distances between probes (③), we gained eigenvectors corresponding to each pair of probes and each PC mode (④). Signs in eigenvectors can be interpreted as the increase/decrease pattern of the distances between probes. (B) A heatmap shows the PCC between the FMND and the frequency of contact between chromosome bins and SNFs (N = 446 cells). Chromosomes colored red, green, and blue correspond to CAF1, CAF2, and CAF3, respectively. \**P* < 0.01 using the Pearson test. (C) Violin plots in the left panel show the PCC between the FMND and the distance from the end of the p-arm. Violin plots in the right panel show the PCC between FMND and the distance from the center of chromosomes. \**P* < 0.01 using the Student’s *t*-test. n.s.: not significant. The X-axis indicates the type of CAF. (D) The bar plot shows the contribution ratio PCA on the correlation matrix in Fig. 5B. (E) The first two PC scores of PCA, which are applied to the correlation matrix in Fig.5B, show the heterogeneity in the association between FMND and the frequency of contact between chromosome bins and SNFs. (F) Bar plots show the first two eigenvectors of PCA on the correlation matrix in Fig. 5B. Bars with different colors mean the different signs in each eigenvector.

Furthermore, we focused on the distribution of MND along the chromosomes by calculating the frequency of the minor direction (FMND) by summing the counts of MND for PCmode1, PCmode2, and PCmode3 along the chromosomes in each chromosomal bin (Fig. S11). Subsequently, an investigation of the correlation between FMND on chromosome bins and the contact frequency with SNFs revealed a correlation matrix, as shown in Fig. 5B. The observed relationships were broadly divided into three categories, depending on the type of chromosome (Fig. 5B). The first category consisted of chromosomes, namely chromosome (Chr) 1, Chr3, Chr5, Chr6, Chr14, Chr17, Chr18, Chr19, and ChrX, where the observed FMND tended to be positively correlated with proximity to markers of constitutive heterochromatin and the nucleolus. This group of chromosomes is designated as chromosomes with anisotropic fluctuation 1 (CAF1). The second category (designated as CAF2) consisted of chromosomes, namely, Chr4, Chr9, Chr10, Chr11, Chr12, and Chr13, where the observed FMND tends to be positively correlated with proximity to a nuclear lamina marker. The third category (designated as CAF3) consisted of chromosomes Chr2, Chr7, Chr8, Chr15, and Chr16, where the observed FMND tended to be positively correlated with the proximity to markers of constitutive heterochromatin, open heterochromatin, nuclear speckles, and transcriptional activity. We then examined the distribution of FMND on the chromosomes in these chromosome groups and found that FMND was significantly and positively correlated with the genomic distance from the p-arm terminus in CAF2 compared with CAF1 and CAF3 (Fig. 5C). This result suggests that the anisotropy of the chromosome fluctuation direction may be affected by the association between the spatial distribution of nuclear compartments in the nucleus and the genomic sequence/epigenetic states along the chromosomes. When performing PCA on the correlation matrix shown in Fig. 5B, it was evident that PC1 and PC2 alone spatially divided the three types of CAFs in this projection plane and accounted for approximately 70% of the correlation matrix (Figs. 5D and 5E). In addition, when the eigenvectors of PC1 and PC2 were examined, the variance of PC1 tended to correspond to heterochromatin/nucleolar and open chromatin/nuclear speckle markers, whereas that of PC2 corresponded to markers of chromatin components and the nuclear lamina (Figs. 5B and 5F).

### Relationship between FMND and repeat DNAs

We analyzed the relationship between FMND and repeat DNAs on chromosomes. Because heterogeneity in the local distribution of repeat DNAs is related to chromosome folding, it has been reported that SINEB1 is relatively abundant in compartment A and LINE-1 in the compartment B (29). We conducted PCA on the correlation matrix of FMND and “occupancy rate of repeat DNAs in chromosomal bins” for each chromosome and described the “heterogeneity” regarding this relationship (Figs. 6A and 6B). By examining the eigenvector of the PC1 (with the largest contribution ratio) that best explains this “heterogeneity” (Fig. 6B), we attempted to search for repeat DNAs associated with this PC. The repeat DNAs examined in this study are shown in Table S3. Fig. 6C shows that repeat DNAs corresponding to outliers of the first eigenvector (see *Materials and Methods*) were identified, which included LINE-L1-L1Md_A, LINE-L1-L1Md_F2, LINE-L1-L1Md_T, LINE-L1-L1-L1VL4, LINE-L1-L1_Mur1, LINE-L1-L1_Mus3, LINE-L1-L1_Mus4, LINE-L1-L1-Lx, and LINE-L1-Lx2. The distribution of the nine LINE-1 subfamilies identified in Fig. 6C and FMND in each chromosome showed a tendency to be negatively correlated in chromosomes where the PC scores of PC1 were negative and positively correlated in chromosomes where the PC scores of PC1 were positive (Figs. 6D and 6E). These results suggest that the relationship between the anisotropy of chromosomal fluctuations and the distribution of repeat DNAs varies among chromosomes. In addition, the PC score of PC1 on the correlation matrix between FMND and “frequency of contact with SNFs” (Figs. 5D and 5E) and the PC score of PC1 on the correlation matrix between FMND and “occupancy of repeat DNAs in chromosomal bins” (Fig. 6B) correlated significantly (PCC: -0.75, Fig. 6E). The association between the occupancy of the nine LINE1 subfamilies (Fig. 6C) in the chromosomal bins and the contact frequency with SNFs showed a similar trend for all nine LINE1 subfamilies (Fig. 6F). These results suggest that the anisotropy of chromosomal fluctuations may be based on the distribution of repeat DNAs, the basis of epigenetic states, on chromosomes and their interactions with nuclear compartments. In contrast, the correlation between the occupancy rate of the nine LINE1 subfamilies and the contact rate with SNFs showed heterogeneity among chromosomes (Fig. S12). This suggests that the interaction patterns between repeat DNAs and nuclear compartments may vary for each chromosome due to differences in the distribution patterns of repeat DNAs and their spatial arrangements within the cell nucleus.

**Figure 6.**
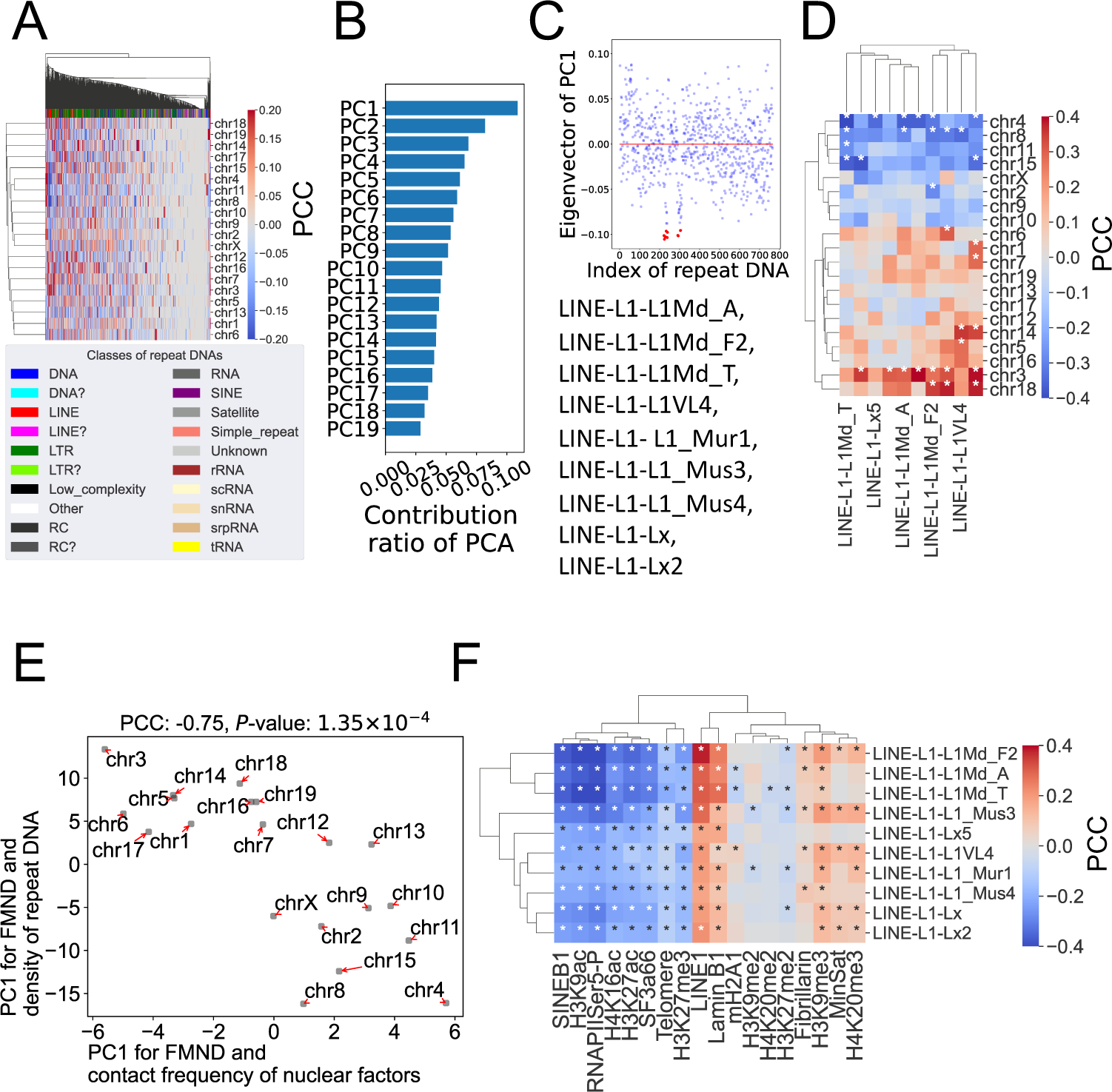
Comparison between FMND along the chromosome and the distribution of repeat DNAs. (A) A heatmap shows the PCC between FMND and the density of repeat DNAs per 1Mb bin (N = 1254 types). Color indexes indicate classes of repeat DNAs (see Table S3). Unknowns and “?” indicate repeat DNAs with unsure classification using the “RepeatMasker” algorithm. (B) A bar plot shows the contribution ratio of PCA on the correlation matrix in Fig. 6A. (C) The eigenvector of PC1 shows the repeat DNAs corresponding to the heterogeneity in the association between FMND along a chromosome and the distribution of repeat DNAs. Red dots show the outliers in the eigenvector of PC1 in Fig. 6B (See Materials and Methods). Types of repeat DNAs shown below the scatter plots correspond to the outliers. (D) A heatmap shows the PCC between FMND and the density of repeat DNAs, which were selected in Fig. 6C, per 1Mb bin (N = 9 types). \**P* < 0.01 using the Pearson test. (E) A scatter plot shows that the X-axis is for the first PC score of PCA on the correlation matrix between FMND and the frequency of contact between chromatin bins and SNFs in Fig. 5E and Y-axis is for the first PC score of PCA on the correlation matrix between FMND and the distribution of repeat DNAs in Fig. 6B (N = 20 types of chromosomes). (F) A heatmap shows the PCC between the contact frequency of SNFs and the density of LINE1 subfamilies for each chromosome bin. \**P* < 0.01 using the Pearson test.

### The relationship between the fluctuation direction of chromosomes and their distribution within the nucleus

We classified chromosomes into three types based on the characteristics of the chromosomal regions associated with anisotropic fluctuations (FMND and contact frequency with SNFs) (Fig. 5B). Subsequently, we investigated the relationship between anisotropic chromosomal fluctuations and the position of chromosomes in a radial distribution within the cell nucleus for each CAF type. Analysis of the relative distance from the nuclear center to the periphery (see *Materials and Methods*) for all cells revealed that the radial distributions within the nucleus for CAF1, CAF2, and CAF3 were identical (Fig. S13A). In the seqFISH+ experiments, as the cell cycle of mESCs was not artificially controlled (25), we investigated whether there were differences in the positional dynamics of CAF related to the cell cycle in terms of their radial distribution within the cell nucleus. To conduct this analysis, we performed PCA on the distance matrix based on the relative distance from the nuclear center to the periphery, as shown in Fig. S13B (Fig. S13C). As shown in Fig. S13D, PCmode1 revealed isotropic fluctuations (contribution rate: 92%) in the direction from the nuclear center to the periphery for all three CAFs. PCmode2 showed fluctuations in which the distribution of CAF3 moved in the opposite direction to that of CAF1 and CAF2 (contribution rate: 5%), and in PCmode3, the distribution of CAF2 moved in the opposite direction to that of CAF1 and CAF3 (contribution rate: 3%) (Figs. S13E and S13F). The contribution of PCmode1, which represented isotropic fluctuations from the nuclear center to the periphery, was 92%. Therefore, the positional dynamics of chromosomes during the cell cycle examined using PCA suggested little differences among the three types of CAF. Furthermore, when we divided the cells into quartile groups based on the PTCF and compared them, no differences in the radial distribution of CAF were observed in any of the quartile groups (Fig. S13G). This suggests that the state of chromosomal condensation/decondensation in individual cells does not lead to differences in the positioning and dynamics of the three types of CAF in the radial distribution within the cell nucleus.

Then, we investigated whether there were differences in the radial distribution of chromosomes between the quartile groups divided by PTCF to reveal the relationship between condensation states of single chromosomes and the radial distribution of chromosomes. We found that the radial distribution of all chromosomes and the three types of CAF were biased toward the center of the nucleus in the highest quartile group (75-100%) of PTCF compared with the lowest quartile group (0-25%) of PTCF (Fig. S13H). In addition, when comparing the Kullback-Leibler divergence, which increased with greater differences between the two probability distributions of the quartile groups, between the quartile group with the highest PTCF (75-100%) and the other quartile groups, the maximum Kullback-Leibler divergence was observed between the highest quartile group (75-100%) and the lowest quartile group (0-25%) for the radial distribution of all chromosomes and the radial distribution of any CAF (Fig. S13I). Furthermore, to elucidate the relationship between anisotropic chromosome fluctuation and the radial distribution of chromosomes, we compared the variance of the PCmode2/PCmode3 PC scores in chromosome fluctuation based on the PTCF quartile groups. Our results showed that for all chromosome types, the variance in PC scores in PCmode2/PCmode3 was higher in the highest quartile group (75-100%) than in the lowest quartile group (0-25%) of PTCF (Fig. S14). In summary, the correlation between anisotropic chromosome fluctuations and the radial distribution of chromosomes suggests that anisotropic chromosome fluctuations tend to occur in cells where chromosomes are positioned near the center of the nucleus.

### Relationship of chromosome fluctuations to interchromosomal condensation/interaction

Finally, we examined the tendency of interchromosomal condensation/interaction within each chromosome group (CAF1, CAF2, and CAF3), considering the correlation between FMND and the contact frequency of SNFs. For this analysis, chromosomal bins were grouped into quartile groups based on FMND, and the means of the interchromosomal distances were examined for each group of chromosomal bins. The means of the interchromosomal distances were shorter in the quartile group (75-100%) with the highest FMND than in the quartile group (0-25%) with the lowest FMND in all CAFs (Fig. 7 left panels).

**Figure 7.**
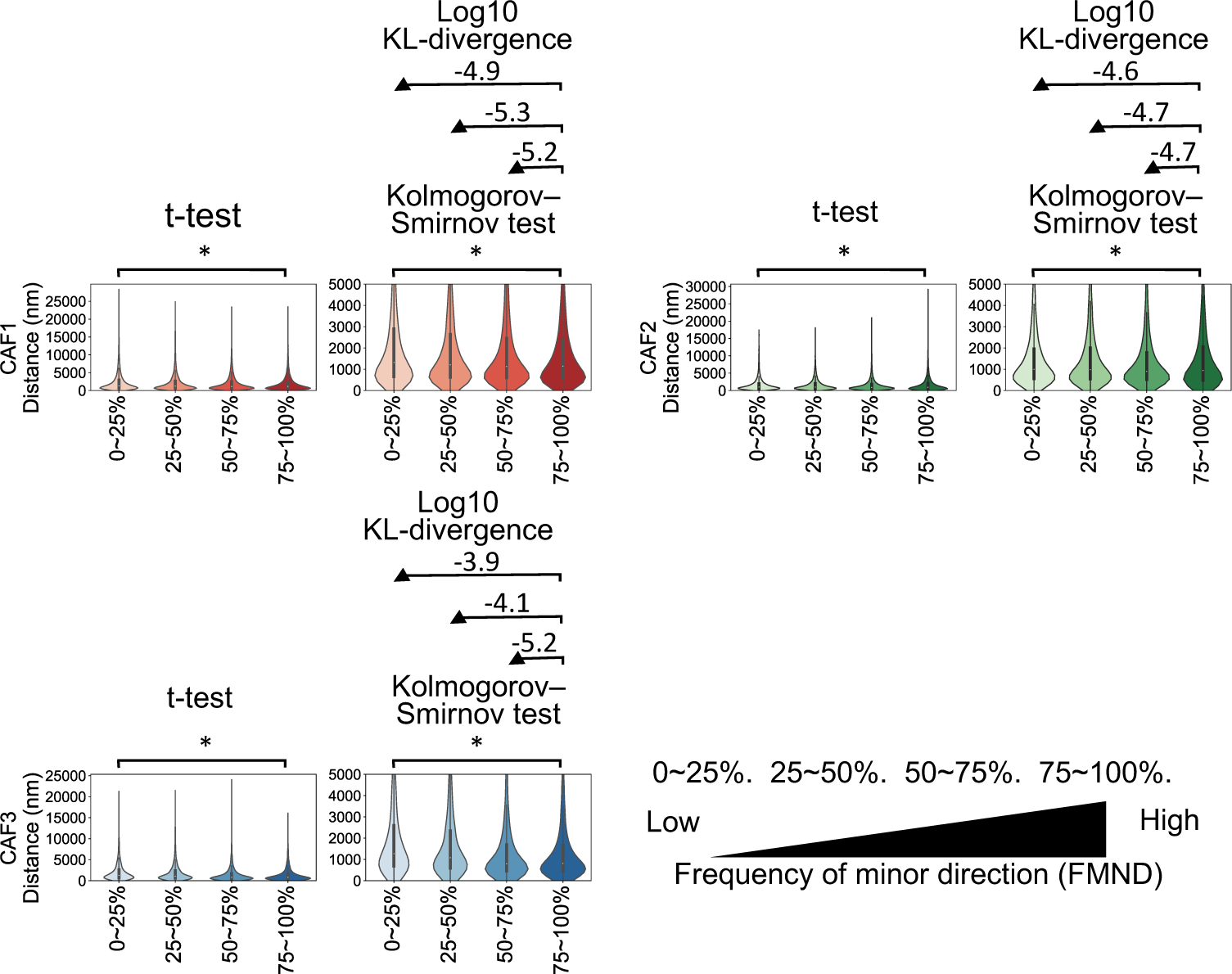
The relationship between FMND and interchromosomal aggregation for each CAF. The violin plot shows that interchromosomal aggregation tends to occur in chromosome bins with anisotropic fluctuations along the chromosomes (N = 446 cells). Each left panel shows the entire distance distribution. Each right panel shows the distance distribution in 0-5000 nm. The red, green and blue panels correspond to CAF1, CAF2 and CAF3, respectively. The X-axis indicates the four groups of chromosome bins equally divided based on the FMND. “75-100%” means the group of chromosome bins with the highest frequency in a minor direction. The Y-axis indicates the interchromosomal distances between each group of chromosome bins. Values above brackets in the left panels indicate the results of the Student’s *t*-test. \**P* < 0.01 using the Student’s *t*-test. n.s.: not significant. Values above the brackets in the right panels indicate the results of the Kolmogorov-Smirnov test. \**P* < 0.01 using the Kolmogorov-Smirnov test. n.s.: not significant. Values above the arrows in the right panels indicate Kullback–Leibler divergence (KL divergence), which measures how one distribution differs from a second.

Next, we examined whether there were differences in the distribution of interchromosomal distances for the four groups of chromosomal bins. Using the Kolmogorov–Smirnov test to assess differences in probability distributions between two quartile groups, we found that, for any CAF, the distribution of interchromosomal distances for the quartile group (75-100%) with the highest FMND was shorter compared with the quartile group (0-25%) with the lowest FMND on the chromosomes (Fig. 7 right panels). In addition, we quantitatively examined the differences between the two probability distributions using the Kullback–Leibler divergence, which increased with greater differences between the two probability distributions of the quartile groups. When comparing the chromosome bin group with the quartile group (75-100%) to the other quartile groups, we found that there was a monotonic increase in Kullback–Leibler divergence as the FMND in the quartile groups of chromosome bins decreased, particularly in CAF3 (Fig. 7 right panels). Furthermore, in other CAFs, the Kullback–Leibler divergence between the quartile group (75-100%) and quartile group (0-25%) was larger compared with the Kullback–Leibler divergence between the quartile group (75-100%) and quartile group (50-75%) (Fig. 7 right panels). These results suggest that interchromosomal condensation/interaction in each CAF is more likely to occur between chromosome regions with relatively high FMND than between regions with relatively low FMND.

## Discussion

### Comparison between PCA-based analysis and polymer simulation method in the chromosome fluctuations

In this study, we investigated the directions of chromosome fluctuations by applying PCA to the 3D data of hundreds of single-chromosome structures based on Euclidean distances obtained using seqFISH+ experiments. The PCA results revealed that PCmode1 was isotropic, while PCmode2 and PCmode3 were anisotropic (Fig. 3C). From our results, we successfully identified, for the first time, that there is anisotropy in chromosome fluctuations using experimental data. However, our observations of the amplitudes of chromosomal fluctuations did not often align with the results of PHi-C2, a polymer simulation based on Hi-C data. This discrepancy is thought to arise from the formulation of PHi-C2’s polymer simulation method, which is formulated as ‘the contact matrix stored in Hi-C data is a function of the variance of the distance distribution between chromosomal domains.’(26, 27).

### Relationship between isotropic/anisotropic chromosome fluctuations and cell cycle

During the transition between the M phase and interphase, all chromosomes undergo chromosomal dynamics through condensation (spindle-like shaped chromosomes) and decondensation (spherical-shaped chromosomes) (30). Therefore, the biological significance of PCmode1 for the fluctuation of chromosome is expected to align with the dynamics related to the transition between the M phase and the interphase because VDMSC, related to the spindle-like/spherical shape of chromosomes, significantly correlates with the PC scores of PCmode1 for the fluctuation of chromosome (Figs. S5, 4A, 4B and S6). In cells with high PTCF values, where chromosomes are often in a condensed state, the polymer-like fluctuations of the chromosomes were observed to be anisotropic (Figs. S9 and S14). This suggests that anisotropic chromosomal fluctuations were observed during the cell cycle process before and/or after the M phase, specifically when chromosomes were spindle-like shape, rather than in the interphase when chromosomes were spherical shape.

The radial distribution of chromosomes in cells from the highest quartile group (75-100%) was biased toward the center of the nucleus compared with the quartile group (0-25%), which was divided based on PTCF (Figs. S13H and S13I). This indicates that there is the fluctuation of chromosome radial distribution that associates with the chromosome condensation and decondensation in the cells sampled as seqFISH+ data. Furthermore, when PCA was applied to the radial distribution of CAFs within the cell nucleus, the obtained PCmode1 was isotropic and had a 92% of contribution ratio (Figs. S13C and S13D). These results suggest that PCmode1 for the fluctuation of chromosome radial distribution also corresponds to with the dynamics related to the transition between the M phase and the interphase.

### The periodicity of anisotropic chromosome fluctuations

Next, we focused on the periods of each PC mode to represent the time scale of chromosomal fluctuations per cycle. The cell cycle time of mESCs is approximately 11 h (31). Herein, we assumed that 11 hours was the period of PCmode1 that corresponds to the behavior associated with the transition between the M phase and the interphase of the cell cycle. Additionally, since the cell cycle was not artificially controlled in the samples of seqFISH+ data (25), it is presumed that single-cell/single-chromosome data of mESCs in seqFISH+ were randomly sampled from this approximately 11-hour cell cycle process. Therefore, the pseudo-timescale between each chromosomal 3D structure/snapshot subjected to PCA in this study was approximately 2 min, considering thar the X chromosome had the smallest number of sampled structures/snapshots (N = 396: Fig. S1E). Assuming that PCmode1 corresponds to the behavior associated with the transition between the M phase and the interphase of the cell cycle, the periods of each PC mode are estimated to fall within approximately 2 min to 11 h. By observing that the PCA eigenvalues are proportional to the square of the period, we calculated the estimated periods for PCmode2 and PCmode3 (Table S4). The estimated periods of PCmode2 and PCmode3 were longer than those of the G1 (62 min), G2 (161 min), and M phases (36 min) but shorter than those of the S phase (355 min) (31). In the future, by using cell cycle control/identification techniques, it is anticipated that applying PCA to single-chromosome samples with different cell cycle phases will enable the analysis of experimental data on chromosome fluctuations specific to each cell cycle phase. For example, it has been reported that replication foci, regions of chromosomes undergoing DNA replication during the S phase, transition from euchromatin in the early stage to heterochromatin in the later stage (32). It is possible to elucidate the correlation between the transition of replication foci on chromosomes during the S phase and chromosomal fluctuations.

### The biological significance of anisotropic chromosome fluctuations

The different correlation patterns between FMND and SNFs by CAF (Fig. 5B) suggest that the interchromosomal space where chromosome fibers with a relatively high FMND anchor may vary for each CAF (Figs. S15A and S15B). Specifically, it has been speculated that in CAF1, it involves nucleolar and heterochromatin bodies, CAF2 involves the nuclear lamina, and CAF3 involves nuclear speckles and heterochromatin bodies (Figs. S15A and S15B). In addition, the maximum contact frequency between the DNA-FISH probes and each SNF was less than 60% (Fig. S16A), suggesting that contacts with nuclear compartments/interchromosomal spaces in any chromosomal region are transient (Figs. S15A and S15C). In previous studies, the chromosome territory-inter-chromosomal space model (CT-IC model) reported that chromosome territories and interchromosomal spaces are not clearly separated; chromosome fibers anchor to the interchromosomal space in the interphase (6-8). Considering the CT-IC model, it is speculated that the relatively high FMND of chromosomal regions is due to the restriction of chromosome fluctuation by the anchoring of chromosome fibers to the interchromosomal space in the interphase (Figs. S15A and S15B). Contrary to this speculation, upon examining the PTCF of cells where contact (< 300 nm) between DNA-FISH probes and SNFs was observed, most probes were found to be in contact with any SNF in cells belonging to the quartile group (50-75%) (Figs. S16B and S16C). Cells corresponding to the quartile group (50-75%) based on PTCF, an index corresponding to the state of chromosome condensation/decondensation in individual cells, are speculated to be in the G2 phase or near the G1 phase around the M phase when chromosomes take on a spindle-like shape. Taken together, it is anticipated that anisotropic chromosome fluctuations occur, especially during the processes around the M phase of the cell cycle, through the interactions between chromosomal regions and nuclear compartments/interchromosomal spaces. Furthermore, nuclear compartments known to contribute to nuclear organization by interacting with specific regions of chromosomes, such as the nucleoli, nuclear speckles, and nuclear lamina, have been reported to disappear by the end of the prophase (33-35). Our results suggest that anisotropic chromosome fluctuations are likely to occur during the processes around cell cycle transitions before and after the M phase (Fig. S14). In summary, these findings imply that anisotropic chromosome fluctuations contribute to the transition of nuclear organization between the interphase, where nuclear compartments appear, and the M phase, where nuclear compartments disappear.

The question of which cell cycle phase, ‘before M phase,’ ‘after M phase,’ or ‘before and after M phase,’ exhibits anisotropic chromosome fluctuation has been speculated based on relevant studies. It has been reported that the asymmetry in chromosome condensation and decondensation occurs during the transition between the interphase and the M phase (36-38). This involves adopting a helical loop array structure and a cylindrical shape with cohesin unloading and condensin loading (36, 37). Conversely, only cohesin loading occurred during the transition from the M phase to the interphase (36, 37). In line with these experimental results, simulation results based on the landscape-switching model using Hi-C data, which depict cell cycle-dependent chromosomal structural phase transitions, showed that the trajectories from the interphase to the M phase and from the M phase to the interphase differed (38). Compared with the transition from the M phase to the interphase, the simulation predicted that the shape of the chromosome was cylindrical in the transition from the interphase to the M phase (38). In this study, we found that the chromosome fluctuation of PCmode2/PCmode3 was anisotropic compared with PCmode1 in the terms of the orientation of chromosomes; this has the potential to make the chromosome structure non-cylindrical (Fig. 3C). Considering these previous reports and our results together, the anisotropic fluctuation of chromosomes observed in this study is expected to occur after the M phase, not before the M phase, and to contribute nuclear reorganization from the M phase to the interphase. To test this hypothesis, we identified the cell cycle phase in which the 3D structure of single chromosomes was bent using DNA-FISH after identifying the cell cycle state of mESCs using immunostaining markers that can identify the G2, prophase, metaphase, anaphase, telophase, and G1 phases.

### The effects of repeat DNAs on the anisotropic fluctuations of chromosomes

Previous studies have reported the exclusive distribution of LINE1 and SINEB1/Alu on chromosomes and that the biased distribution of these repeat DNAs roughly corresponds to the position of TAD formation (29). Additionally, LINE1 is closely associated with nuclear organization, and the spatial localization of LINE1 corresponds to the spatial distribution of the nucleolar and nuclear lamina (7, 13, 29, 39). In this study, we identified nine types of LINE1 elements that contributed the most to the direction of PC1, the principal variance, in the correlation matrix between FMND and repeat DNAs. When examined across chromosomes 1-19 and X, the density of these nine LINE1 types was found to have a consistent pattern of correlation with contact frequency with any SNF (Fig. 6F). However, when examined on a chromosome-by-chromosome basis, the observed correlation varied among chromosomes (Fig. S12). This suggests that owing to differences in the distribution patterns of repeat DNAs and the associated epigenetic states on each chromosome, the roles of the same types or families of repeat DNAs in nuclear organization may differ. In the future, it will be important to investigate the relationship between the distribution patterns of repeat DNAs on chromosomes and the topology of chromosome distribution in correspondence with the distribution of nuclear compartments to understand the conditions under which the transient anchoring of chromosome fibers to the interchromosomal space occurs.

## Materials and Methods

### PHi-C2 analysis using HiC-seq data

The read counts from the pre-processing of the HiC-data were trimmed using Trimgalore (https://github.com/FelixKrueger/TrimGalore#readme) with default settings. HiCUP v0.8.2 (40) with Bowtie2 v 2.3.4.1 (41) was used to map and filter di-tags to the mouse genome to construct the GRCm38. To estimate complex compliance along each chromosome, we performed PHi-C2 analysis (27) on Hi-C data with a 1Mb chromosomal bin using default settings.

### seqFISH+ data processing

#### Identification of single chromosomes

To consider chromosome fluctuations as a polymer, it is necessary to divide sequential DNA-FISH signals into single chromosomes. To avoid dealing with signals from multiple chromosomes as much as possible as those from a single chromosome, we designed an algorithm to identify single chromosomes as follows:

1^st^: Get signals of seqFISH+ (25) from the same types of chromosomes in each cell nucleus.

2^nd^: Find the expected minimum of signals per single chromosomes using a binomial distribution. The “binom.isf” function in scipy (https://scipy.org/about/) was used, where q (the survival function residual), p (the probability of a single success), and N (the number of samples) were set at 0.9999, 0.5 that is the detection efficiency of seqFISH+ (25) and the number of designed probes per chromosome.

3^rd^: Find the median of the distances between the signals in single chromosomes. First, we calculated the distances between the signals in the same types of chromosomes for all combinations. Second, we find the median from the lower half of the distance in ascending order. This step was modified to find the median from all distances for the chromosome X because the seqFISH+ data were from the male cell line.

4^th^: Perform Density-Based Spatial Clustering of Applications with Noise (DBSCAN) from scikit-learn (https://scikit-learn.org/stable/), where “min_samples” and “eps” were set as the expected minimum of signals per single chromosomes calculated in the 2^nd^ step and the median of distances between signals in single chromosomes calculated in the 3^rd^ step.

5^th^: Remove the clusters defined in the 4^th^ step suspected of containing signals from multiple chromosomes. The anticipated maximum of signals per single chromosome from seqFISH+ data was determined using the inverse survival function of the binomial distribution, employing the “binom.isf” function in scipy (https://scipy.org/about/). In this calculation, the parameters “q,” “p,” and “n” were assigned values of 0.0001, the reciprocal of 2 (representing the number of identical autosomes in the nucleus), and the count of signals from the same chromosomes in the seqFISH+ data, respectively. This step was omitted for chromosome X.

#### Preprocessing and clustering analysis for fluorescent signals

To determine the distribution of the nuclear compartments, we focused on the following fluorescent signals of nuclear factors (SNFs): RNAPIISer5P, H3K9ac, H3K27ac, H4K16ac, H3pSer10, H3K9me2, H3K27me2, H3K27me3, H4K20me1, H3K9me3, H4K20me2, H4K20me3, SF3a66, Fibrillarin, mH2A1, LaminB1, SINEB1, LINE1, MinSat, and Telomere (Table S1). The signal intensity of each biomarker was processed to obtain a z-score for each nucleus. Voxels with a z-score less than zero were removed to consider informative nuclear compartments in the seqFISH+ dataset. A GMM from scikit-learn (https://scikit-learn.org/stable/) was used to divide the voxels into an arbitrary number of clusters. To determine reasonable GMM cluster number parameters between 2 and 60 clusters, the BIC was checked for each trial. The cluster parameter of the GMM was set to 60 because the BIC gradually decreased with an increasing number of clusters (Fig. S3A). Each cluster divided by GMM was annotated based on the criteria in Table S2. The following nuclear compartments were annotated: repeat elements, nuclear speckles, nucleoli, constitutive heterochromatin, transcriptionally active compartment-high, transcriptionally mixed compartment, and transcriptionally active compartment-low.

#### Analysis of the fluctuation amplitude and eigenvector along chromosome PCA

To describe the fluctuation amplitude and eigenvector along the chromosome, we performed a PCA against the pairwise distance matrix of single chromosomes, where the coordinates of the pairwise distance matrix were pairs of chromosome bins. Herein, we used the PCA algorithm that allows missing values, “pca-magic” (https://github.com/allentran/pca-magic). After the first PCA conduction, we removed single chromosomes that correspond to the outliers in the first three PC scores from following analysis on chromosome fluctuation as polymers (see Statistical analyses). Subsequently, we performed the second PCA conduction. The eigenvalues and eigenvectors were interpreted as the fluctuation amplitude and direction along the chromosome, respectively. The IFA based on the coordination of the pairwise distance matrix was calculated as follows:

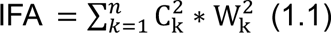

where *C*, *W*, and *n* are the eigenvalues, eigenvectors, and number of PC modes in the PCA, respectively. Note that the radial frequency variable ω of chromosome fluctuation in this calculation is assumed to be constant.

#### Quantification of anisotropy in each eigenvector based on the orientation of the chromosome

To describe the anisotropy of each eigenvector based on the orientation of the chromosome, we applied the following method: First, we constructed a matrix with eigenvectors based on the coordinates of the pairwise distance matrix (original matrix, Fig. 3B, left panels). Next, we flipped the original matrix vertically and horizontally to obtain a flipped matrix (Fig. 3B, middle panels). Third, we calculated the Pearson correlation coefficient (PCC) between the original and the flipped matrices as the index of anisotropy in each eigenvector based on the orientation of the chromosome.

#### Definition of MND for chromosome fluctuation and quantifying the FMND for each chromosome bin

To describe the tendency within direction of each PC mode, we defined chromosome bins with MND. Eigenvectors of the PCA against the pairwise distance matrices of single chromosomes can be interpreted as the direction of chromosome fluctuation in each PC mode. Herein, we defined the MND as the elements with fewer signs in each of the first three eigenvectors. To quantify the FMND for each chromosome bin, we summed the number of MND for the chromosome bins for the first three PC modes.

#### Spatial embedding of eigenvector along chromosomes into mean structures of chromosomes

To visualize the direction of fluctuation along the chromosome, we focused on the eigenvectors of the PCA against the pairwise distance matrices of single chromosomes. First, we construct mean 3D chromosome structures by applying multidimensional scaling to the mean of the pairwise distance matrices for each type of chromosomes. Second, the unit vectors for each pair of chromosomal bins were found in the constructed mean 3D chromosome structures. Third, we embedded the first three eigenvectors of PCA against the pairwise distance matrices of single chromosomes by multiplying each eigenvector by the unit vectors corresponding to pairs of chromosome bins. Finally, we combined the vectors, eigenvectors multiplied by the unit vectors corresponding to pairs of chromosome bins for each chromosome bin, for each chromosome bin.

#### Examining the influence of the fluctuation of one PC mode on the fluctuation of another PC mode

We set the *x*th PC mode as the effector and the *y*th PC mode as the follower. To examine the influence of the effector PC mode on the follower PC mode, we divided the PC scores of the follower as:

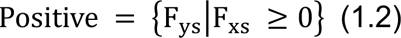

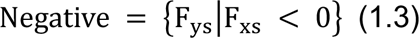

, where F*_ks_*, *k*, and *s* are the matrices of the PC scores, *k*th PC mode, and the *s*th sample, respectively. Next, we compared the variances between *Positive* and *Negative* as follows:

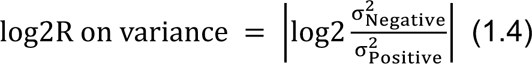

where σ^2^ is the variance of corresponding sets. “*log2R on variance*” can be interpreted as the index of how much the fluctuation of the effector PC mode suppresses the fluctuation of the follower PC mode.

#### Analysis of the shape of single-chromosome structures

To compare the shapes of single-chromosome structures, we performed PCA on the 3D distribution of DNA-FISH probes for each single chromosome. The contribution ratio of the first three PCs provided information about the 3D structure as the ratio of the length of the longest side to that of the two sides perpendicular to it.

#### Estimation of cell trajectory based on the VDMSC for each type of chromosomes

To understand the state of cells based on the variance among DNA-FISH probes from single chromosomes, we performed PCA on the matrix of the mean VDMSC of the same chromosome types for each cell. We selected the PC scores of PC1 as the PTCF.

#### Estimation of the periods of each PC mode based on the actual time

Based on the isotropy of the first eigenvector (Figs. 3C, S4, and S9), we interpreted the fluctuations in the PCmode1 as corresponding to the condensation and divergence of chromosomes related to the cell cycle. The *k*th eigenvalue can be represented as:

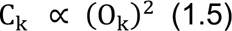

where O*_k_* is the *k*th relative period. Therefore, O*_k_* can be expressed as follows:

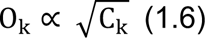

We estimated the periods of the *k*th PC mode (T*_k_*) from (1.5) as follows:

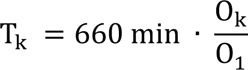

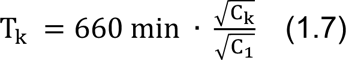

, where 660 min is the doubling time of approximately 11 h for mESCs (31).

#### Measurement of relative distance from the center of the nucleus for CAFs

To describe the relative distribution of each CAF from the center of the nucleus, we calculated the median relative distance between the elements of each CAF and the center of the nucleus as follows. The ‘center_of_mass’ function (https://scipy.org/about/) determined the coordinates of the center of the nucleus in each cell based on the signals of DAPI. To measure the relative distributions of the probes based on radial distribution, the following functions were used:

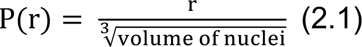

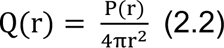

where r, P(r), and Q(r) represent the Euclidean distance from the nuclear center to the periphery, relative distances scaled with the volume of nuclei, and relative distances scaled with the theoretical probability distribution of probes, which is the surface area of the sphere. We used the median Q(r) for the relative distribution of CAFs as the representative values for single nuclei.

#### PCA for the fluctuation of positioning for each type of chromosome from the center to the periphery of the nucleus

To elucidate the fluctuation of radial distribution of each CAF from the center to the periphery of the nucleus, we applied PCA (pca-magic: https://github.com/allentran/pca-magic) to the relative distance matrix from the center of the nucleus, which consisted of the median of each CAF. The eigenvectors from the PCA were interpreted as the direction of fluctuation for each chromosome type.

### Statistical analyses

Correlation analysis was performed using the ‘pearsonr’ function in scipy (https://scipy.org/about/). Specifically, we conducted a Pearson test between the IFA along chromosomes and the complex compliance along chromosomes simulated with PHi-C2, assuming ω of chromosome fluctuation tested using PCA on seqFISH+ as constant. Student’s *t*-test was conducted using the ‘ttest_ind’ function with ‘equal_var’ set to False in scipy (https://scipy.org/about/). Kolmogorov–Smirnov test was conducted using the ‘kstest’ function in scipy (https://scipy.org/about/). PCA was performed using ‘pca-magic’ (https://github.com/allentran/pca-magic). Outliers were detected by checking the survival function of the chi-square distribution generated by squaring the z-scores of the continuous one-dimensional probability distributions (the threshold: < 0.01).

## Data availability

Hi-C data for mouse ESCs were downloaded from the Gene Expression Omnibus (Accession number: GSE96107). seqFISH+ data were downloaded from the Zenodo software (https://zenodo.org/record/3735329). Nested repeat elements on mouse reference genome mm10 were downloaded from the UCSC Genome Browser (https://hgdownload.soe.ucsc.edu/goldenPath/mm10/database/nestedRepeats.txt.gz). Scripts for this study are available at https://doi.org/10.5281/zenodo.10602901.

## Acknowledgements

Computers were partially performed on the National Institute of Genetics (NIG) supercomputer at Research Organization of Information and Systems NIG, Japan. This work was partially supported by JST, the establishment of university fellowships towards the creation of science technology innovation, Grant Number JPMJFS2129.

## Author Contributions

T.N., H.T., A.A., and Y.K. designed the research plan; H.T., A.A., and Y.K. supervised the research; T.N. performed bioinformatics analysis; T.N. and Y.K. wrote the manuscript; T.N., H.T., A.A., and Y.K. finalized the manuscript. All authors have read and agreed to the published version of the manuscript.

---

**Fig. S1.**
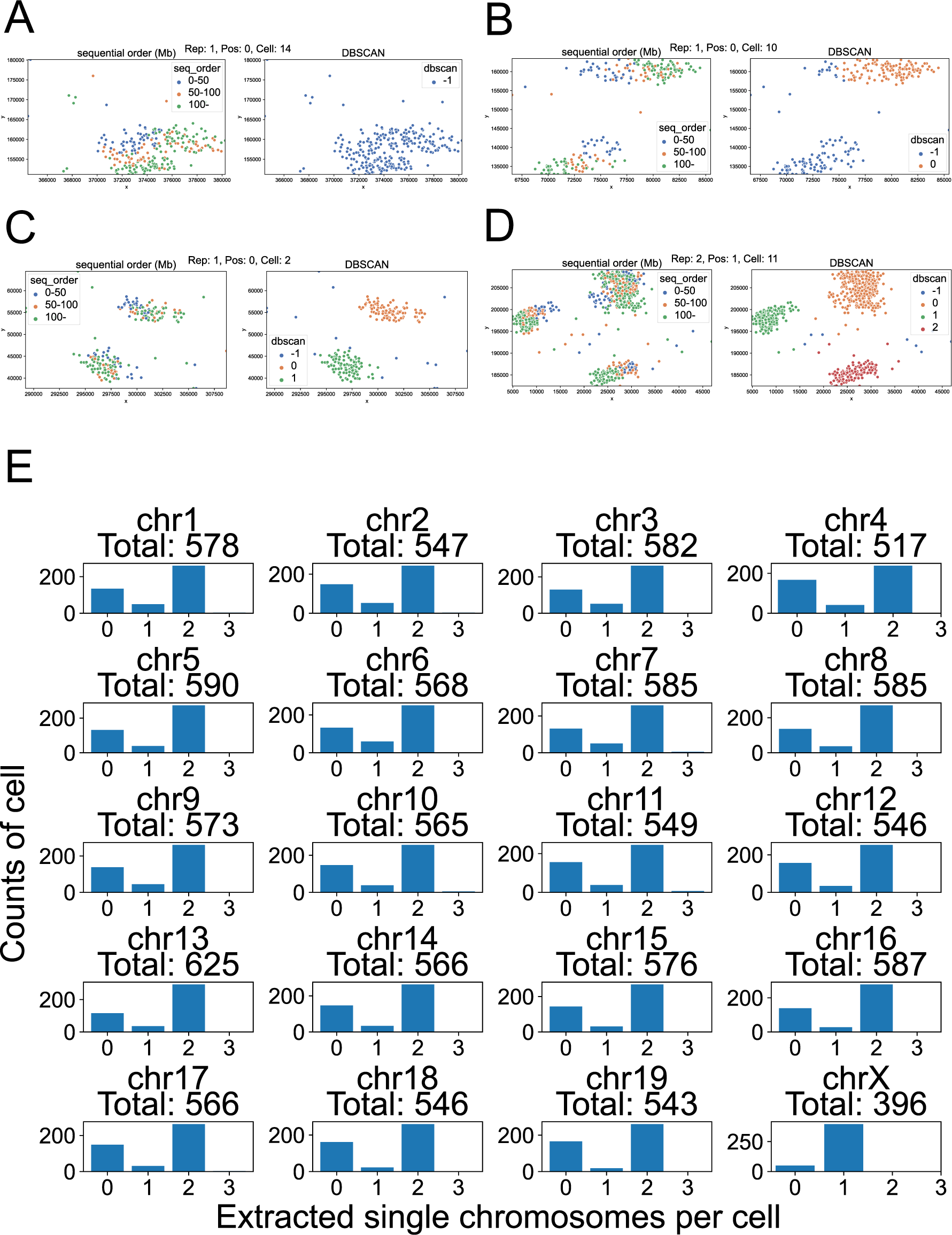
Identification of a single chromosome for each cell nucleus. (A-D) Examples of each single chromosome 1 identified in this study. The left panels show the sequential order of probes on the linear genome using three colors (blue: 0-50Mb, orange: 50-100Mb, green: 100Mb-). The right panels show the single chromosome 1 estimated using DBSCAN (-1: noisy DNA-FISH signals, the other indexes: DNA-FISH signals from single chromosome 1). (E) The number of single chromosomes per cell that were extracted in this study (N = 446 cells). Values next to “Total” mean the total single chromosomes extracted in this study.

**Fig. S2.**
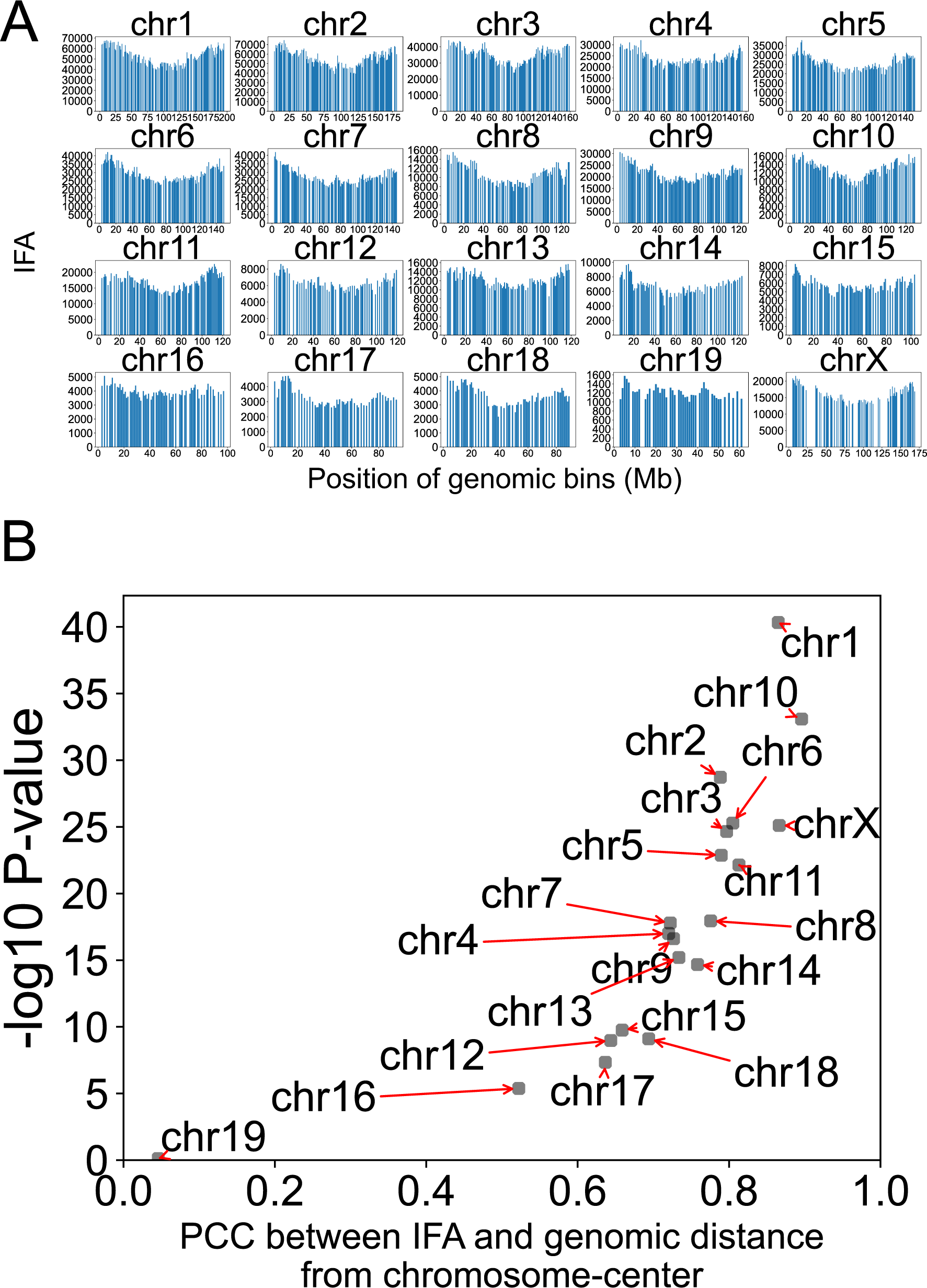
IFA of chromosomes is higher in the ends of chromosomes. (A) The index of fluctuation amplitude (IFA), calculated from chromosome structures, along chr 1 (N = 578), 2 (N = 547), 3 (N = 582), 4 (N = 517), 5 (N = 590), 6 (N = 568), 7 (N = 585), 8 (N = 585), 9 (N = 573), 10 (N = 565), 11 (N = 549), 12 (N = 546), 13 (N = 625), 14 (N = 566), 15 (N = 576), 16 (N = 585), 17 (N = 566), 18 (N = 546), 19 (N = 543), and X (N = 396). Missing values are chromosome bins without designed DNA-FISH probes. (B) A scatter plot shows the correlation between the IFA and the genomic distance from the center of chromosomes. X- and Y-axes indicate the coefficients and *P*-values of the Pearson test.

**Fig. S3.**
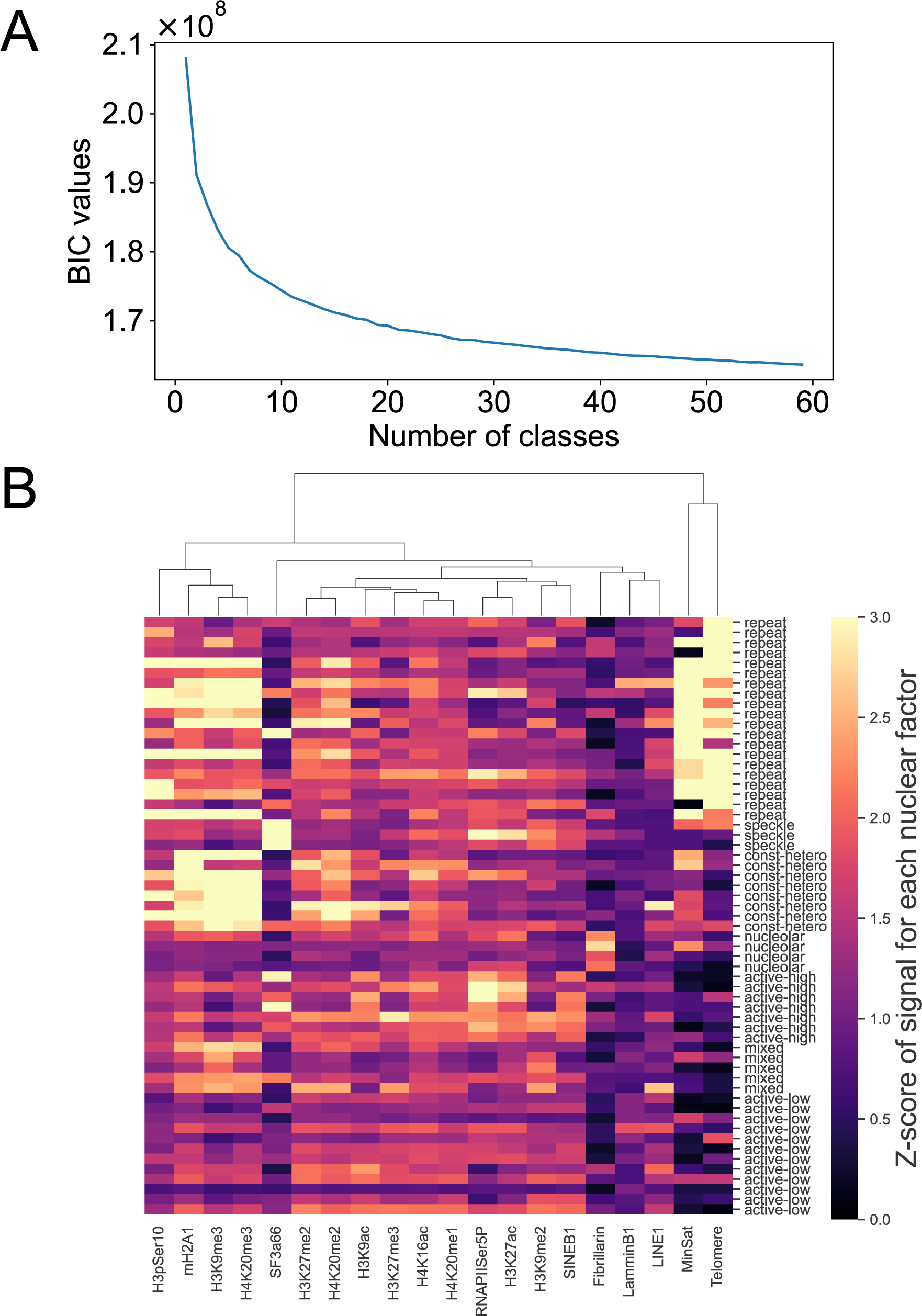
Identification of nuclear compartments. (A) Bayesian information criterion (BIC) of GMM for clustering nuclear compartment based on signals of biomolecules from seqFISH+. The X-axis shows the number of clusters for the GMM parameter. The Y-axis shows BIC values corresponding to the number of clusters. (B) The properties of nuclear compartments annotated using GMM set the number of clusters as 60. A color bar indicates the z-score of the intensity of SNFs from seqFISH+.

**Fig. S4.**
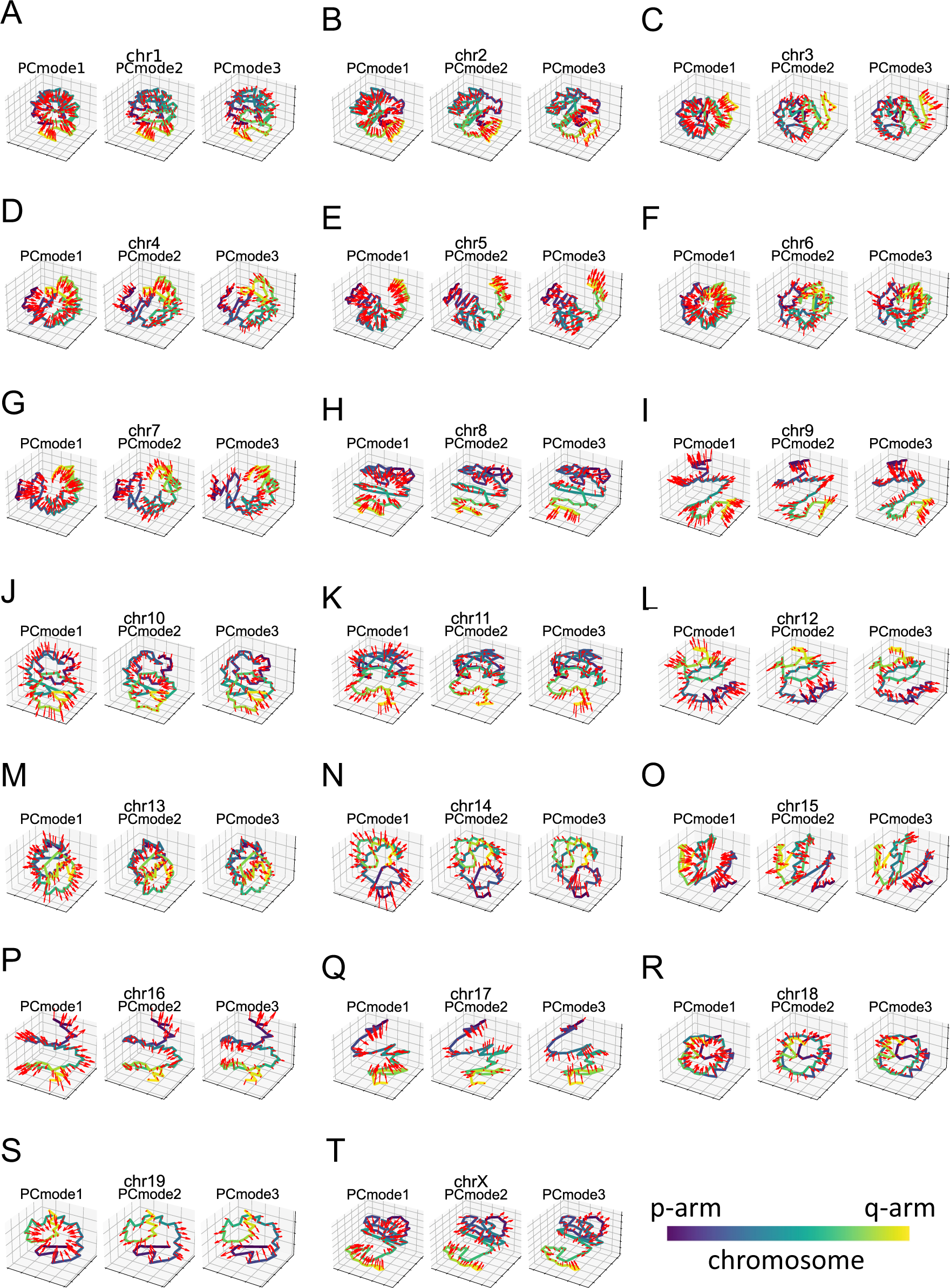
Visualization of the direction of chromosome fluctuation. (A-T) Red arrows show the direction of chromosome fluctuation in the first three PC modes in the mean structures of chr 1 (N = 578), 2 (N = 547), 3 (N = 582), 4 (N = 517), 5 (N = 590), 6 (N = 568), 7 (N = 585), 8 (N = 585), 9 (N = 573), 10 (N = 565), 11 (N = 549), 12 (N = 546), 13 (N = 625), 14 (N = 566), 15 (N = 576), 16 (N = 585), 17 (N = 566), 18 (N = 546), 19 (N = 543), and X (N = 396). A color bar shows the relative position along chromosomes (i.e., The darkness of the color scale means the proximity from the end of the p-arm).

**Fig. S5.**
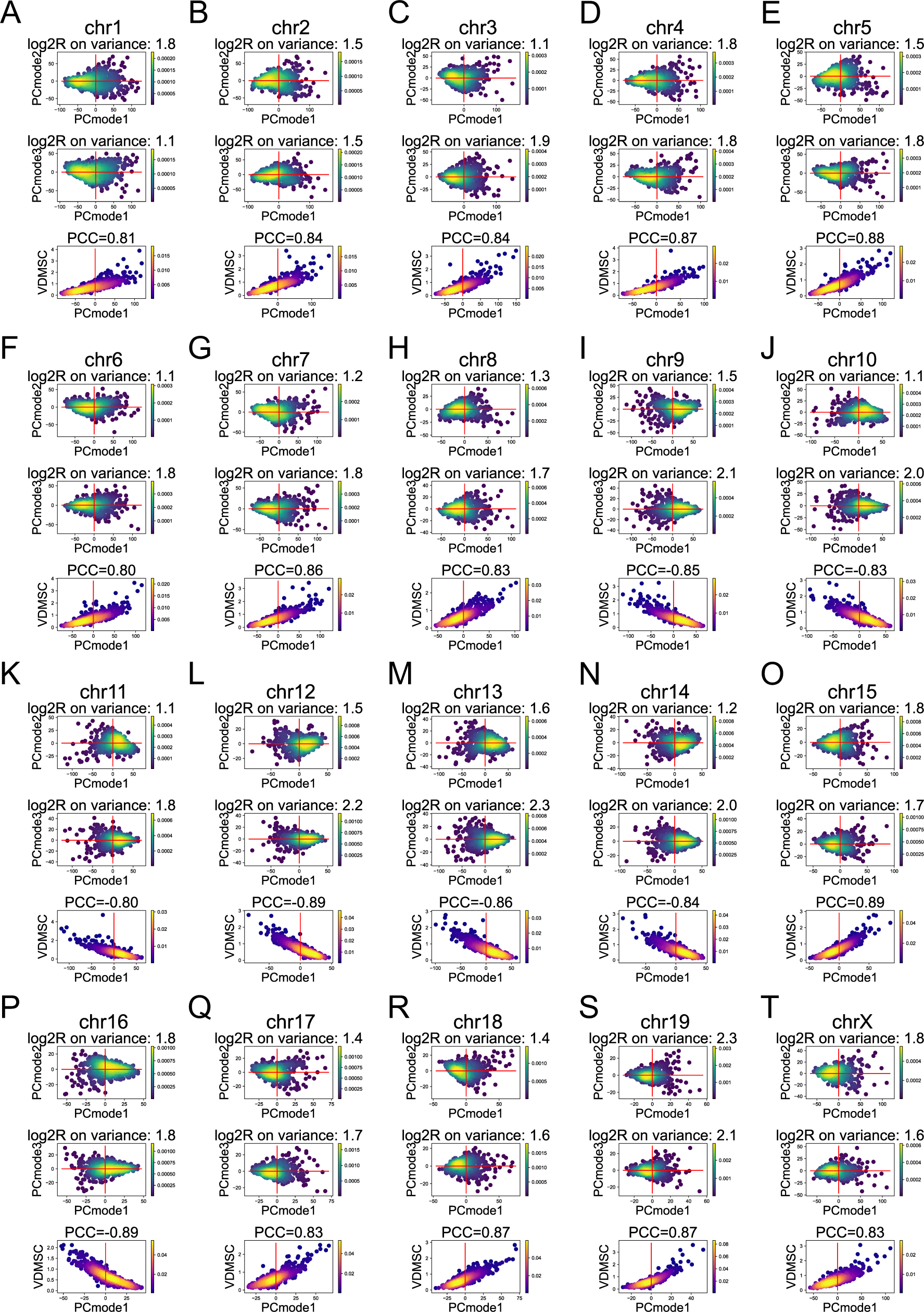
The probability distribution in the PC scores of the first three PC modes. (A-T) The upper and middle panels show the probability distribution in the PC scores of the first three PC modes for chr 1 (N = 578), 2 (N = 547), 3 (N = 582), 4 (N = 517), 5 (N = 590), 6 (N = 568), 7 (N = 585), 8 (N = 585), 9 (N = 573), 10 (N = 565), 11 (N = 549), 12 (N = 546), 13 (N = 625), 14 (N = 566), 15 (N = 576), 16 (N = 585), 17 (N = 566), 18 (N = 546), 19 (N = 543), and X (N = 396). Red lines show the 0 in each PC mode. log2R on variance means the absolute log2 ratio between variances of PC mode corresponding to the Y-axis based on negative and positive samples of PC mode corresponding to the X-axis. Lower panels show the probability distribution in the relationship between PCmode1 and variances of distance matrix for single chromosomes.

**Fig. S6.**
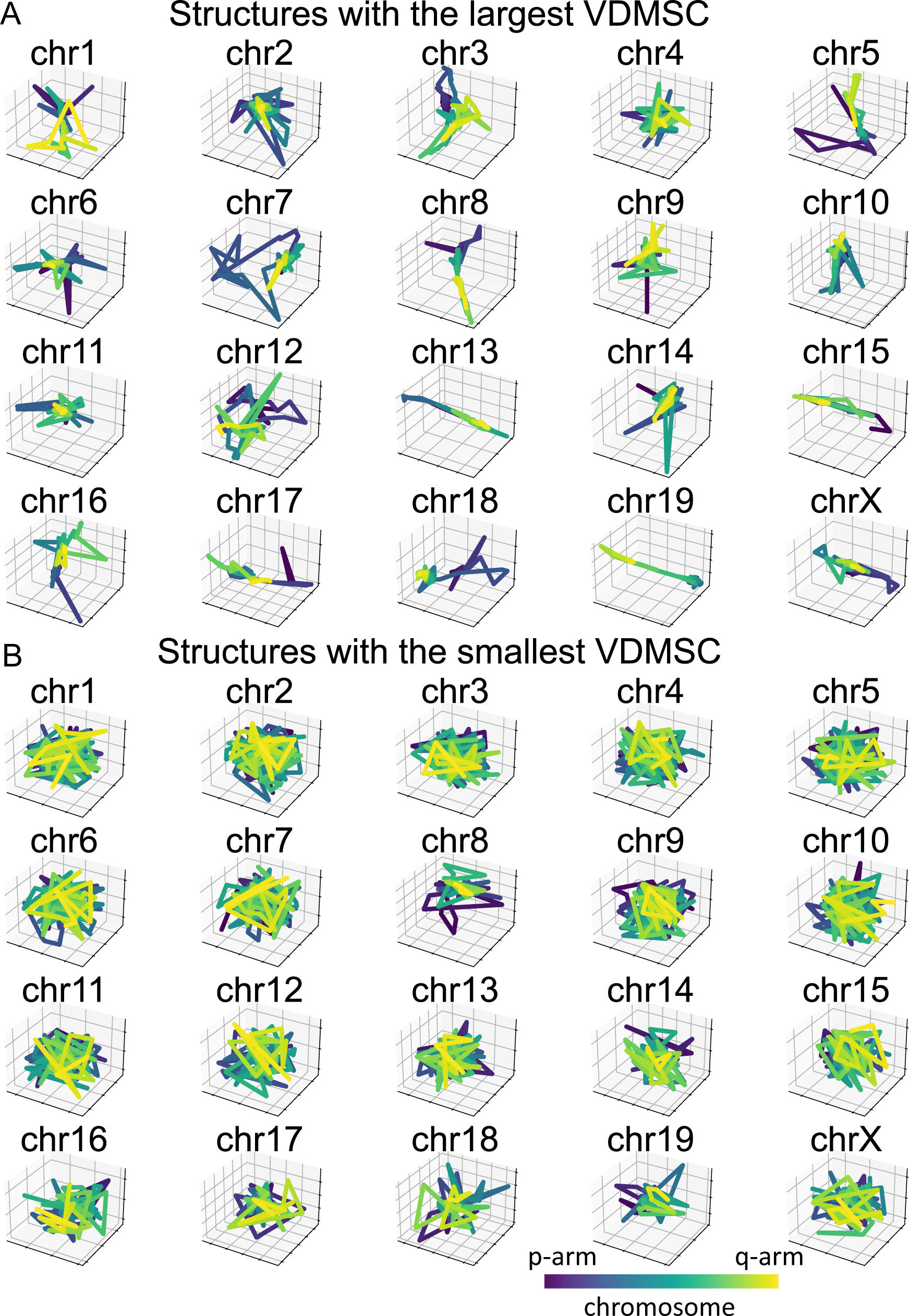
The representative structures of chromosomes correspond to VDMSC. (A and B) Structures of chromosomes 1-19 and X with the largest (A) and smallest (B) VDMSC. A color bar shows the relative position along chromosomes (i.e., The darkness of the color scale means the proximity from the end of the p-arm).

**Fig. S7.**
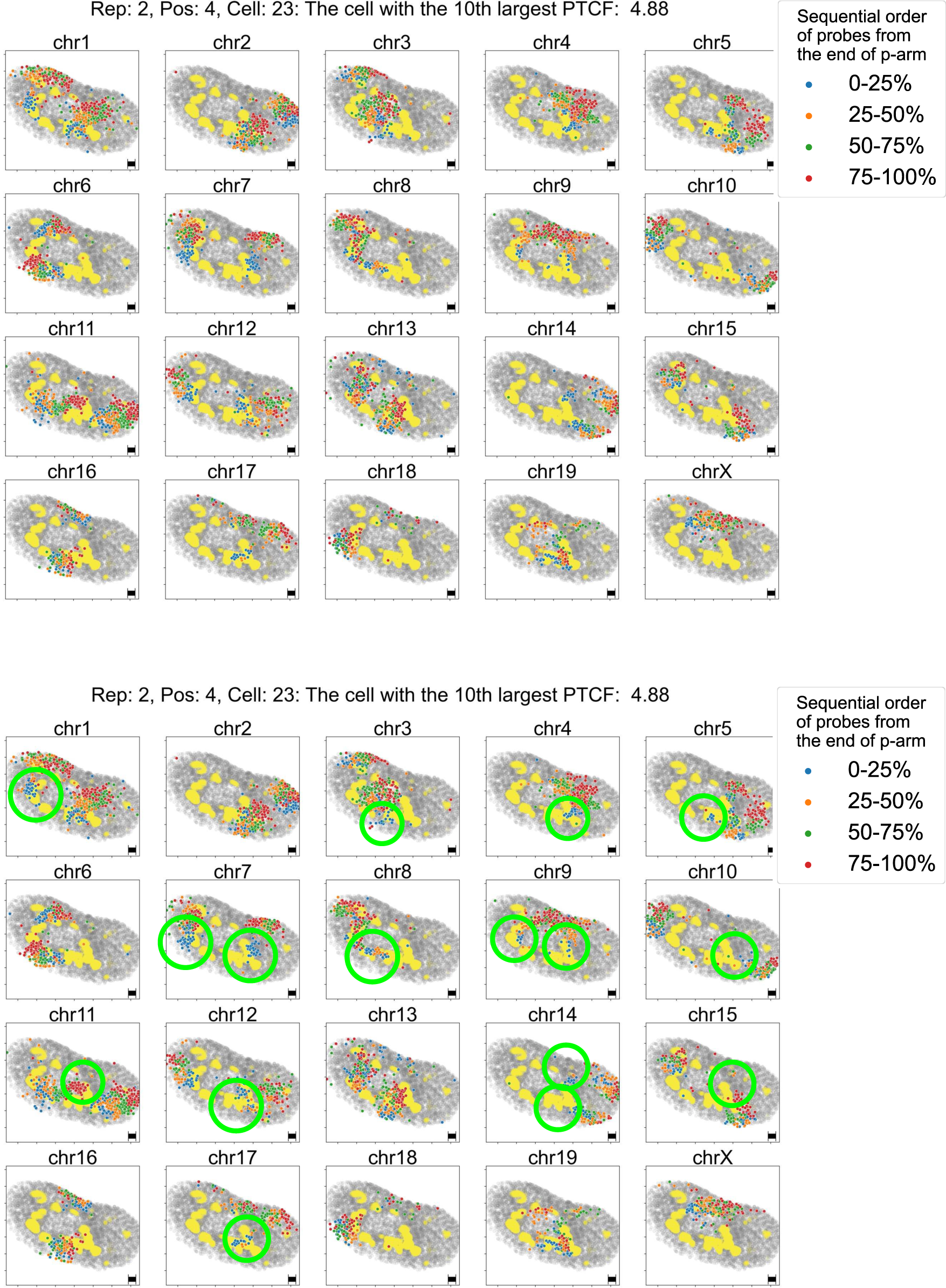
Chromosome structures in the representative cell correspond to the 10th largest PTCF value. Chromosomal structures in cells with the 10th largest PTCF. The blue, orange, green, and red dots correspond to the genomic coordination of probes at 0-25%, 25-50%, 50-75%, and 75-100% from the end of the p-arm. The gray dots indicate the positions of all probes without the chromosomes of interest. The yellow dots indicate the positions of fibrillarin, which is a nucleolar marker. The green circles in the lower panel indicate the chromosome regions corresponding to the anisotropic parts of the chromosome structures. Black bars indicate 1µm for X- and Y-axes in seqFISH+ data.

**Fig. S8.**
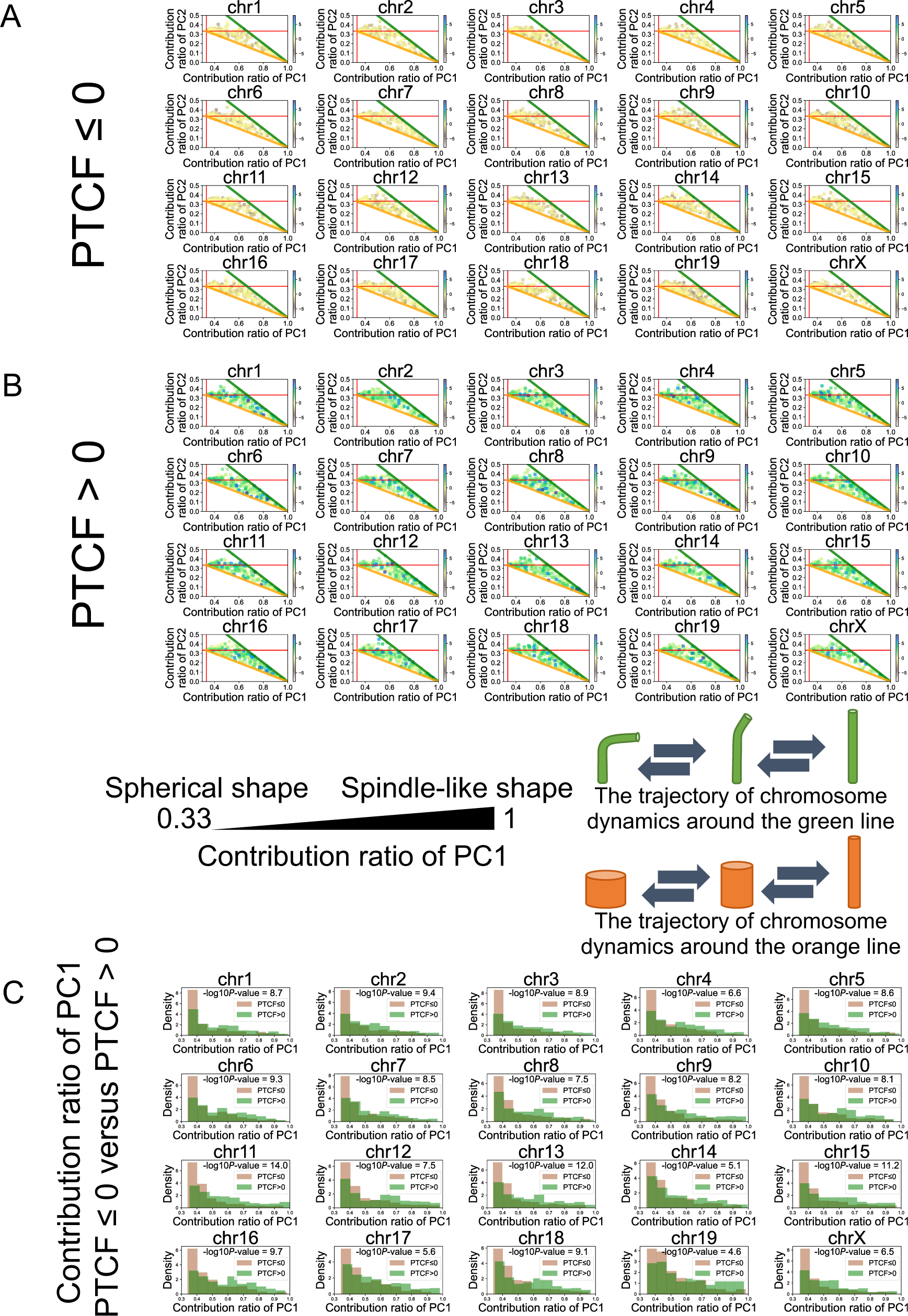
The shape of single chromosomes for each type of chromosome. (A and B) Scatter plots show the contribution ratio of PC1 and PC2 of PCA on the 3D distribution of probes from each single chromosome. The contribution ratio of PC1 can be interpreted as the index for the shape of single chromosomes from a spherical to a spindle-like state. Each dot indicates the shape-based state of the single chromosome for Chr 1 (N = 578), 2 (N = 547), 3 (N = 582), 4 (N = 517), 5 (N = 590), 6 (N = 568), 7 (N = 585), 8 (N = 585), 9 (N = 573), 10 (N = 565), 11 (N = 549), 12 (N = 546), 13 (N = 625), 14 (N = 566), 15 (N = 576), 16 (N = 585), 17 (N = 566), 18 (N = 546), 19 (N = 543), and X (N = 396). Color bars indicate the PTCF of the cell corresponding to each single chromosome. Chromosome dynamics around green lines correspond to the bending of spindle-shaped single chromosomes. Chromosome dynamics around orange lines correspond to the expansion and contraction of spindle-shaped single chromosomes. (C) Density-based histograms show the distribution of the contribution ratio of PC1 from single-chromosome structures from positive and negative PTCF cells. -log10*P*-values indicate the results of the Kolmogorov–Smirnov test.

**Fig. S9.**
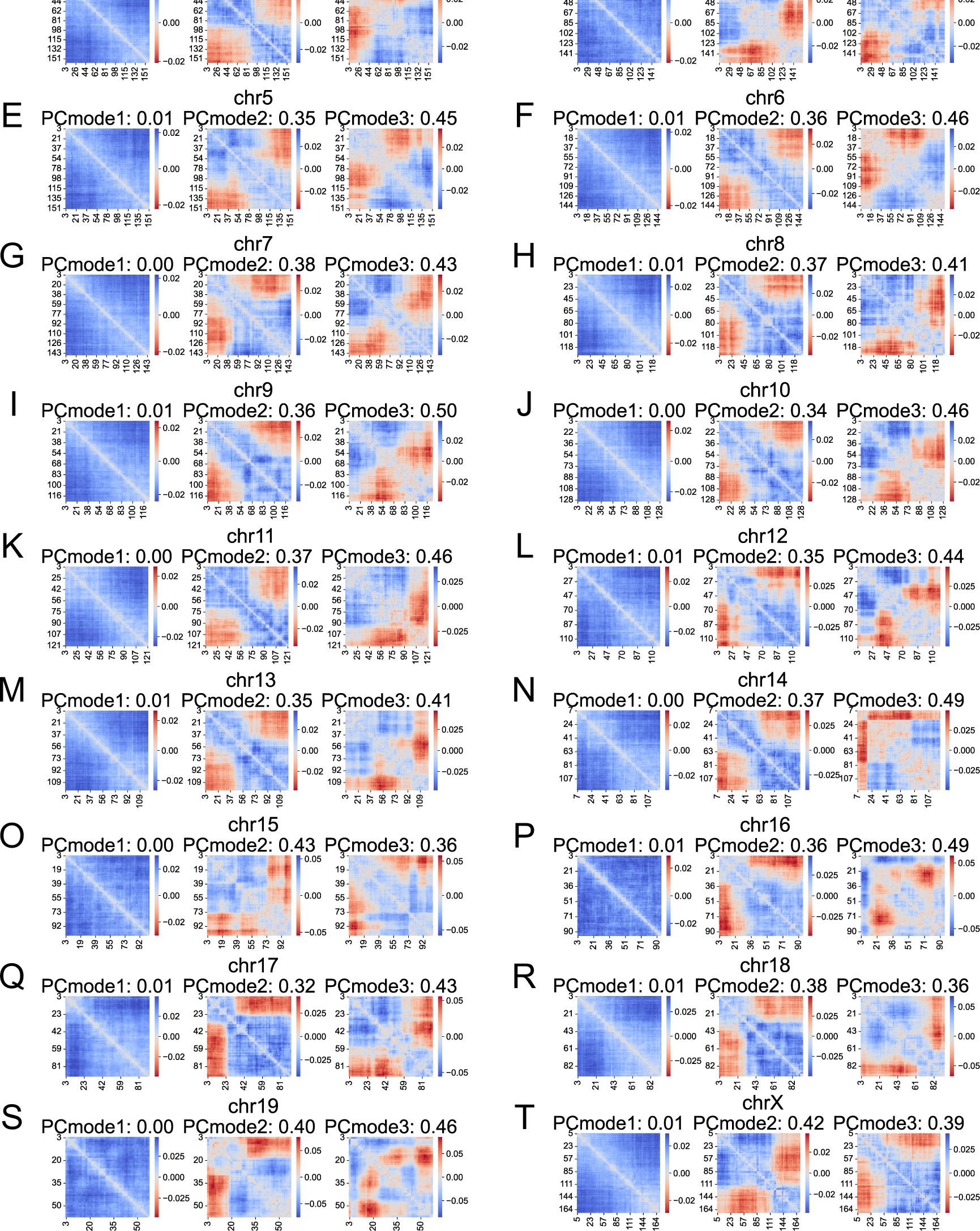
The anisotropic fluctuation along the chromosome for the first three PC modes. (A-T) The eigenvector of the first three PC modes in Chr 1 (N = 578), 2 (N = 547), 3 (N = 582), 4 (N = 517), 5 (N = 590), 6 (N = 568), 7 (N = 585), 8 (N = 585), 9 (N = 573), 10 (N = 565), 11 (N = 549), 12 (N = 546), 13 (N = 625), 14 (N = 566), 15 (N = 576), 16 (N = 585), 17 (N = 566), 18 (N = 546), 19 (N = 543), and X (N = 396). X- and Y-axes indicate the genomic coordinate of the chromosomes (Mb). The signs of eigenvectors with red color indicate the minor direction (MND) in each PC mode. The values in each panel indicate the ratio of pairs of bins, which are in the MND for each eigenvector.

**Fig. S10.**
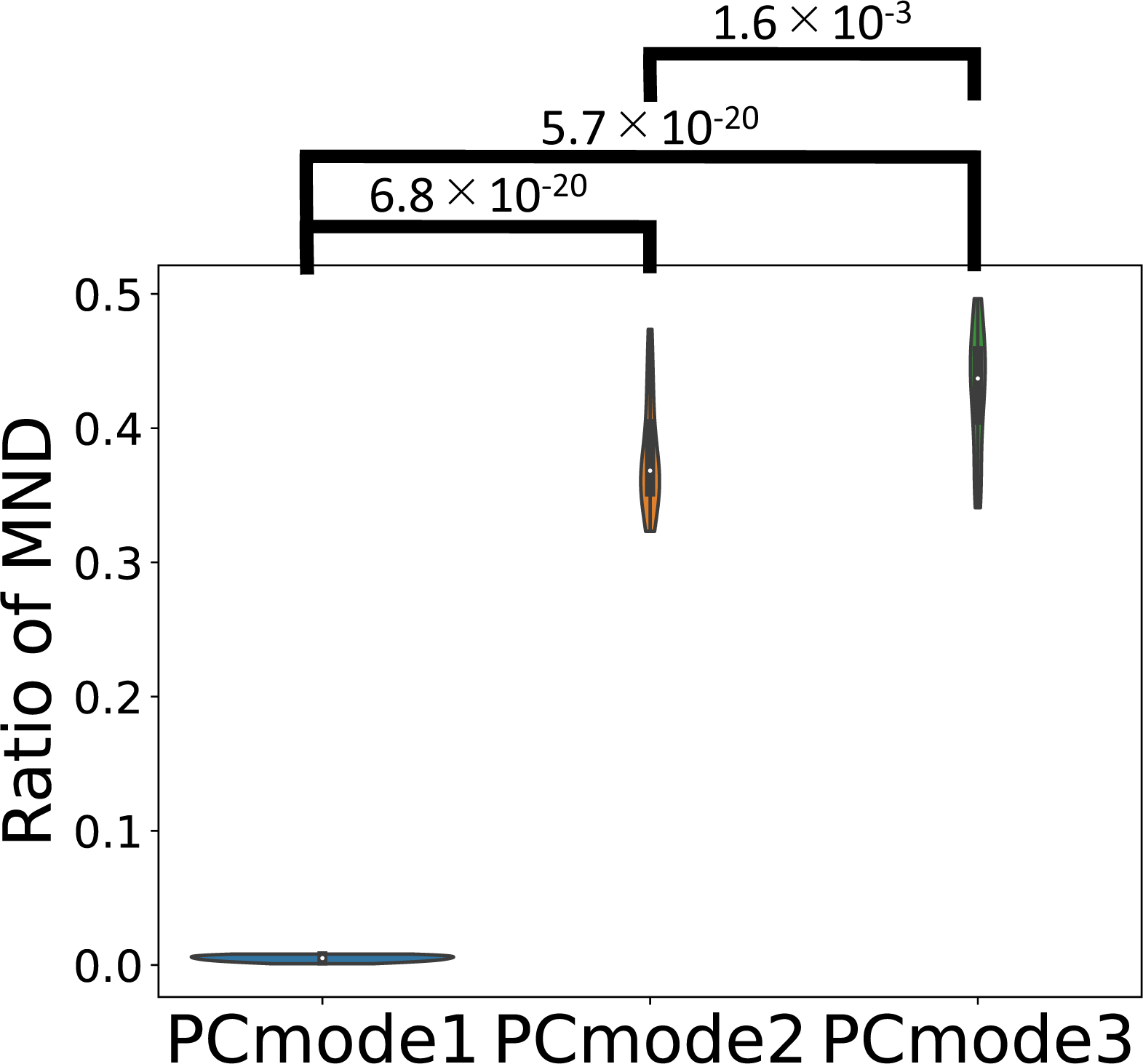
**The ratio of MND in the first three PC modes.**The violin plot shows the ratio of the pairs of bins in the MND for each eigenvector. Values indicate the results of the Student’s *t*-test as *P*-value.

**Fig. S11.**
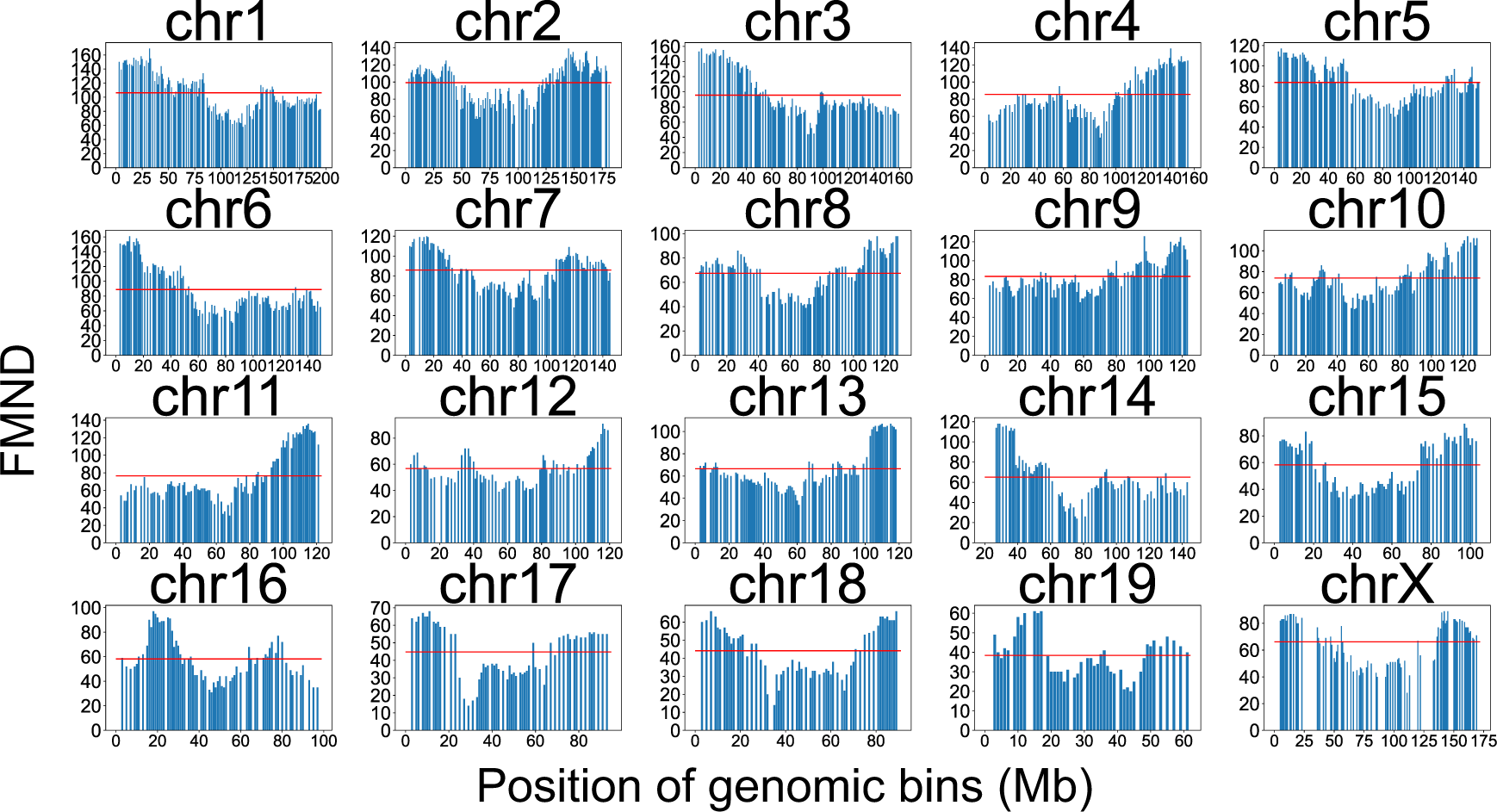
The frequency of anisotropic chromosome fluctuation in the first three PC modes. The total number of the MND in each chromosome bin, FMND, the first three PC modes along Chr 1 (N = 578), 2 (N = 547), 3 (N = 582), 4 (N = 517), 5 (N = 590), 6 (N = 568), 7 (N = 585), 8 (N = 585), 9 (N = 573), 10 (N = 565), 11 (N = 549), 12 (N = 546), 13 (N = 625), 14 (N = 566), 15 (N = 576), 16 (N = 585), 17 (N = 566), 18 (N = 546), 19 (N = 543), and X (N = 396). Red lines show that the mean of the FMND. Missing values are chromosome bins without designed DNA-FISH probes.

**Fig. S12.**
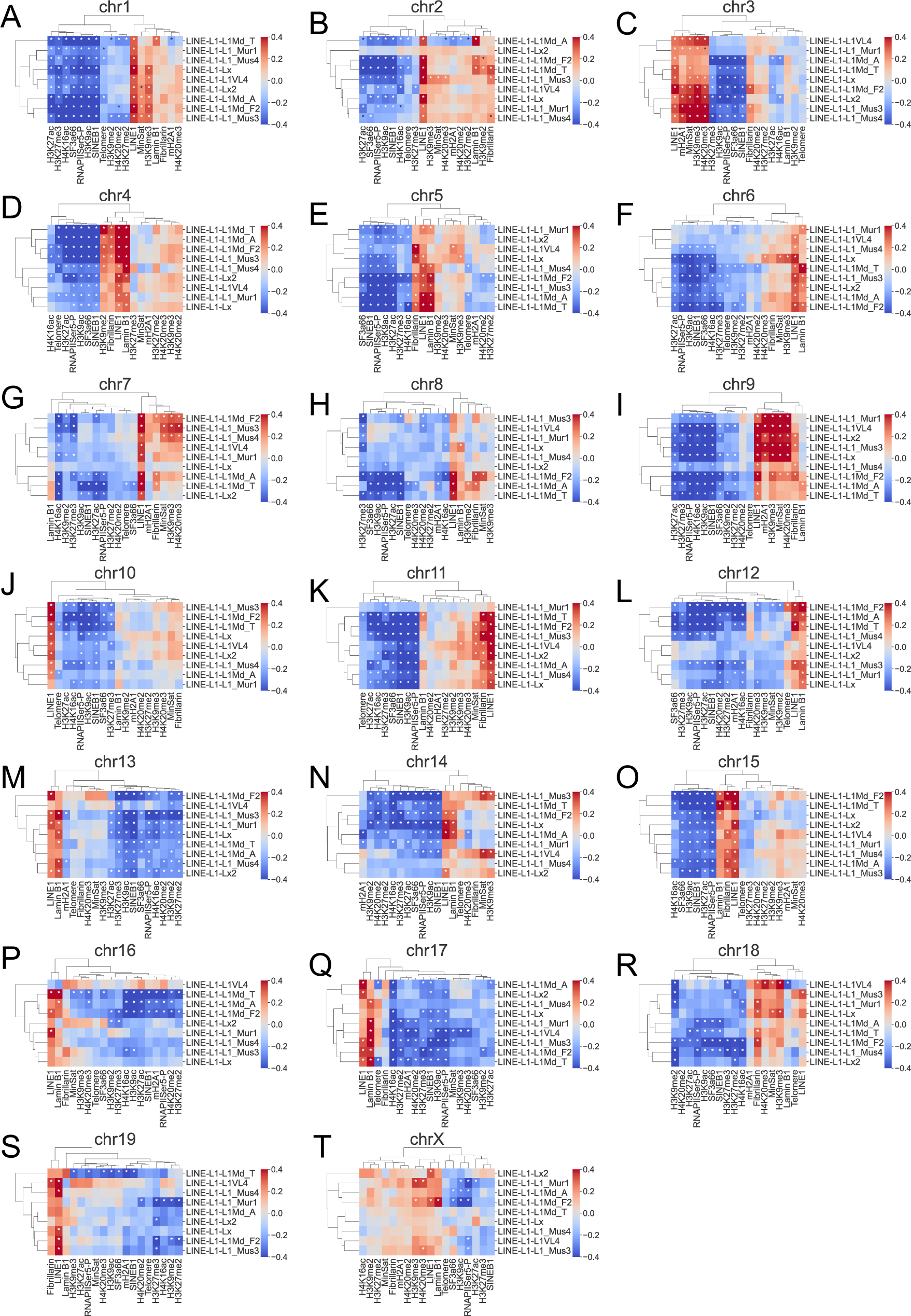
The association between the contact frequency of SNFs and the density of LINE1 subfamilies. (A-T) Heatmap shows the PCC between the contact frequency of SNFs and the density of LINE1 subfamilies for Chr 1 (A: N = 178 bins), 2 (B: N = 185 bins), 3 (C: N = 152 bins), 4 (D: N = 149 bins), 5 (E: N =140 bins), 6 (F: N = 154 bins), 7 (G: N = 142 bins), 8 (H: N = 113 bins), 9 (I: N = 134 bins), 10 (J: N = 130 bins), 11 (K: N = 125 bins), 12 (L: N = 107 bins), 13 (M: N = 119 bins), 14 (N: N = 100 bins), 15 (O: N = 100 bins), 16 (P: N = 91 bins), 17 (Q: N = 83 bins), 18 (R: N = 82 bins), 19 (S: N = 65 bins), and X (T: N = 120 bins). \**P* < 0.01 using the Pearson test.

**Fig. S13.**
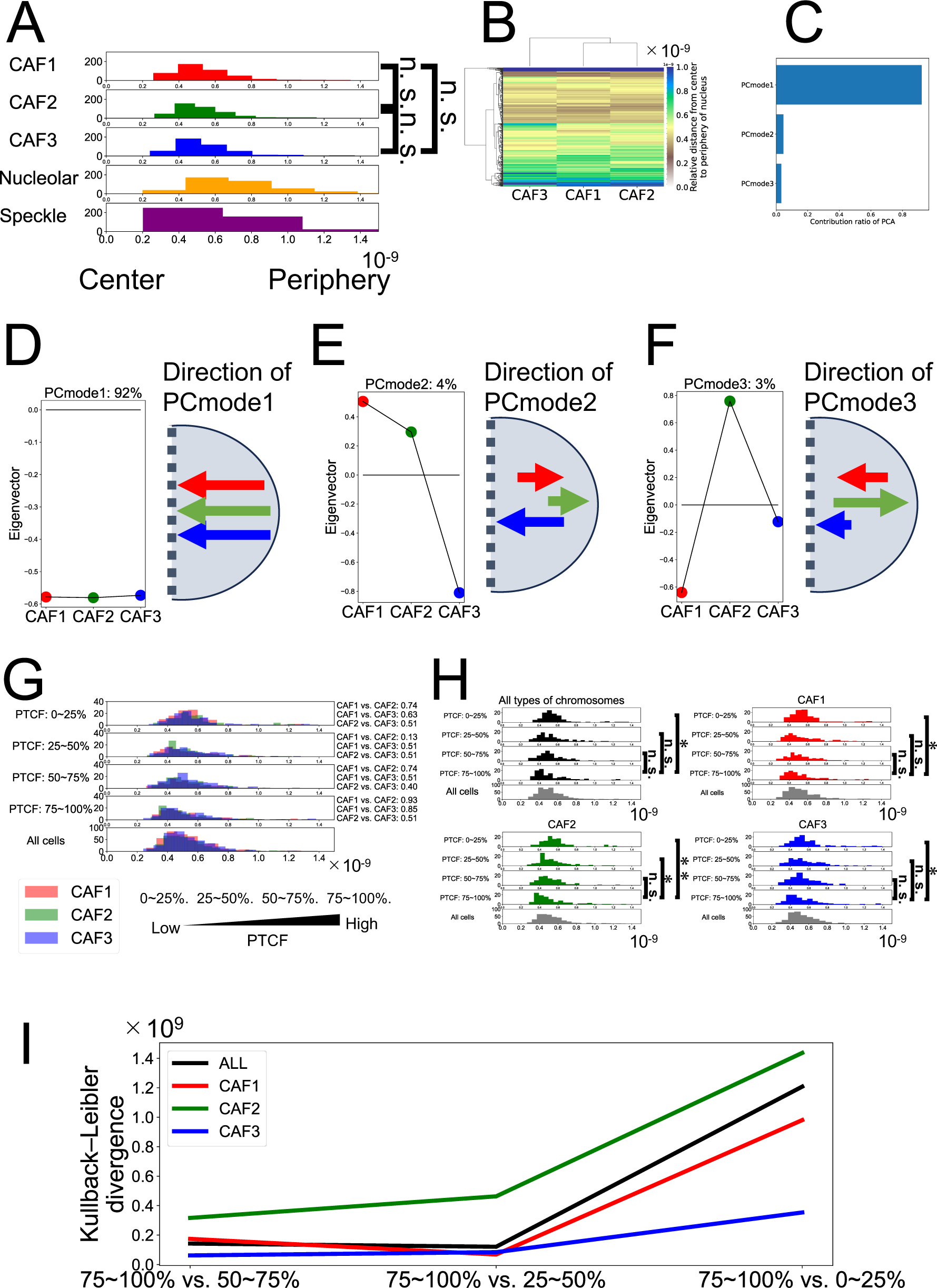
The distribution and fluctuation of positioning for each type of chromosomes along center to periphery of nuclear. (A) Histograms show that the range of positioning for each type of CAF in Fig. 5B (N = 446 cells). X-axis indicates that relative position along center to periphery of nuclear. The right panel shows the results of Kolmogorov-Smirnov test, a test for the equality of two continuous one-dimensional probability distributions. n.s.: not significant by Kolmogorov-Smirnov test (P ≥ 0.01). (B) A heatmap shows the median of the relative distance matrix from the center of the nuclear for each type of CAF in Fig. 5B (N = 446 cells). (C) A bar plot shows the contribution ratio of PCA applied to the relative distance matrix from the center of the nucleus in Fig. S13B. (D-E) The first three eigenvectors from PCA applied on the relative distance matrix from the center of the nucleus in Fig. S13B show that fluctuation direction for each type of CAF from the center to the periphery of the nucleus. (G) Histograms show the radial distribution of each CAF in the cell corresponding to the quartile groups based on PTCF (N = 427 cells). The values on the right side of each panel indicate the result of Kolmogorov-Smirnov test on the two radial distributions of CAFs in the corresponding quartile group as *P*-value. X-axis indicates that relative position along center to periphery of nuclear. (H) Histograms show that the range of chromosome positioning in each quartile group of cells corresponding to PTCF (N = 427 cells). The right panels show the results of the Kolmogorov-Smirnov test. *: *P* < 0.05 using the Kolmogorov-Smirnov test. **: *P* < 0.01 using the Kolmogorov-Smirnov test. n.s.: not significant. (I) Kullback-Leibler divergence of the range of chromosome positioning between the PTCF related quartile group “75∼100%” and the other quartile groups.

**Fig. S14.**
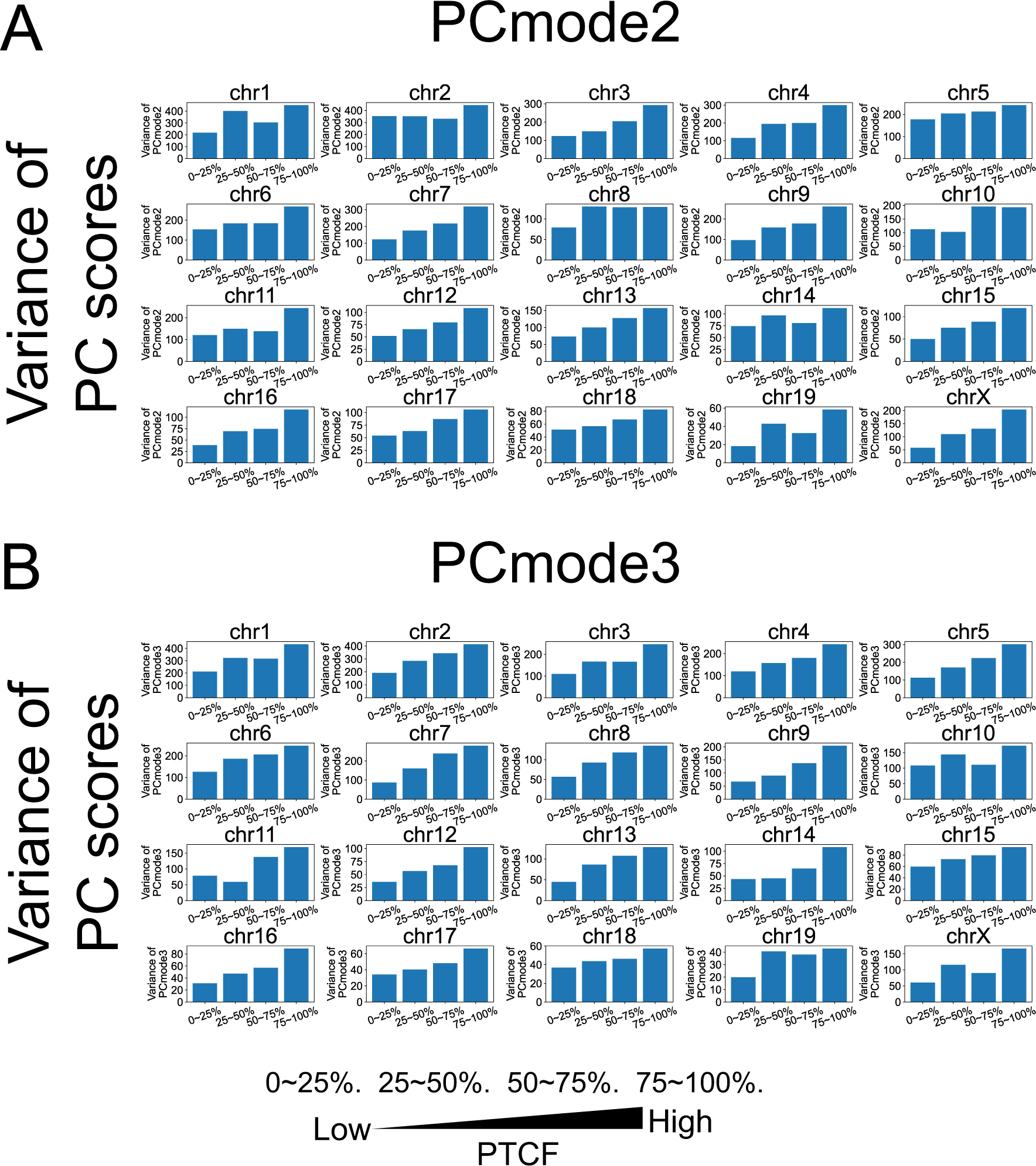
The association between PTCF and the variances of PC scores in PCmode2/PCmode3 for the chromosome fluctuation. (A and B) Comparison of the variances of PC scores in PCmode2 (A)/PCmode3 (B) of chromosome fluctuation among quartile groups of PTCF.

**Fig. S15.**
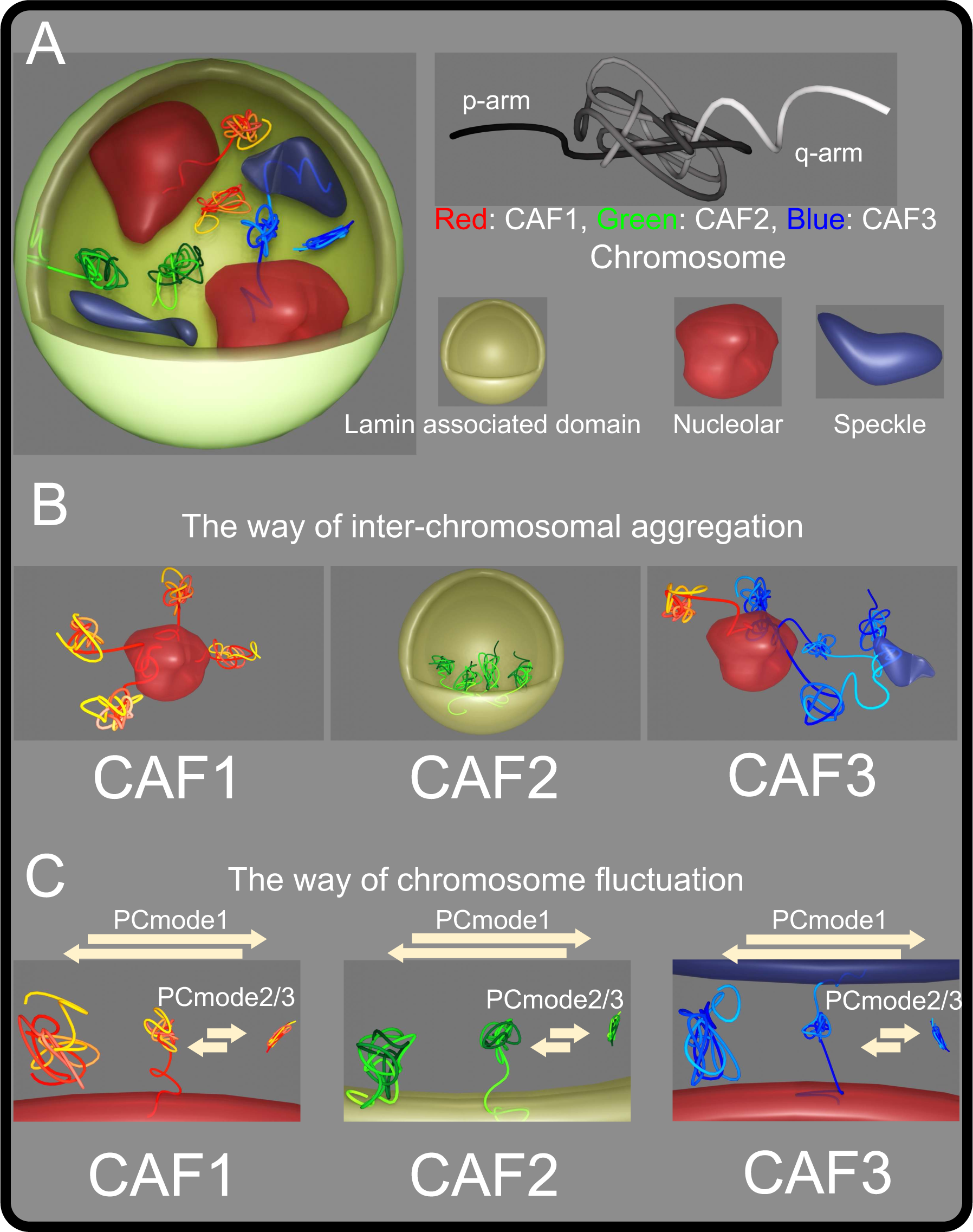
The model on the chromosome fluctuation in the nucleus that contains nuclear compartments/structures. (A) The trend in the association between chromosome fluctuation and the positioning of chromosomes against nuclear compartments. Chromosome regions with pale colors indicate the direction of q-arm and vice versa. (B) The interchromosomal aggregation of each CAF in mESCs. (C) The way of chromosome fluctuation of each CAF in mESCs.

**Fig. S16.**
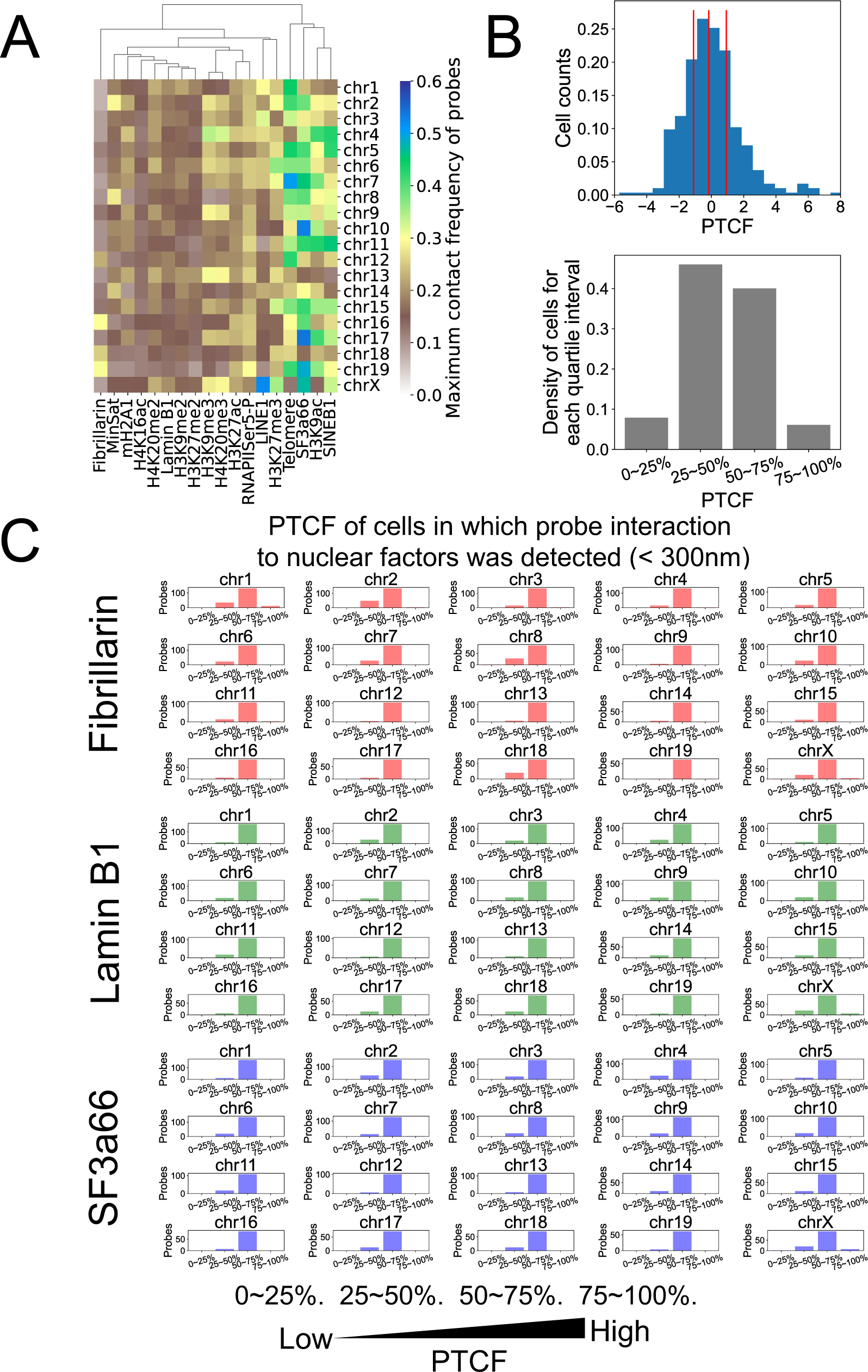
The property of contact between probes and SNFs. (A) A heatmap shows the maximum contact frequency of probes for each chromosome. Contact frequencies of probes were calculated as the ratio of cells whether a probe colocalizes with a SNF within 300nm. (B) The density of cell for each quartile interval based on PTCF. Red lines indicate each quartile. (C) PTCF quartiles to which the median PTCF of cells in which each probe interacts with fibrillarin (< 300 nm) belongs. The X-axis is the count of probes. The Y-axis is the quartile groups of PTCF.

**Table S1.** The SNFs that were considered in this study.

| Serial number | Name of nuclear factors | Abbreviation of nuclear factors | Position in nuclear | Description of nuclear factors |
| --- | --- | --- | --- | --- |
| 1 | Telomere | - | Genomic DNA | A marker of the ends of chromosome |
| 2 | MisSat | Minor satellite | Genomic DNA | A marker of pericentromere |
| 3 | SINEB1 | Short interspersed nuclear elements B1 | Genomic DNA | - |
| 4 | LINE1 | Long interspersed nuclear elements 1 | Genomic DNA | - |
| 5 | mH2A1 | Histone variant macroH2A 1 | Chromatin | A marker of constitutive-heterochromatin |
| 6 | H3K9ac | Acetylation at the 9th lysine residue of the histone H3 | Chromatin | A marker of open chromatin |
| 7 | H3K9me2 | Di-methylation at H3K9 | Chromatin | A marker of transcription inactivation |
| 8 | H3K9me3 | Tri-methylation at H3K9 | Chromatin | A marker of constitutive-heterochromatin |
| 9 | H3K27ac | - | Chromatin | A marker of open chromatin |
| 10 | H3K27me2 | - | Chromatin | A marker of transcription inactivation |
| 11 | H3K27me3 | - | Chromatin | A marker of transcription inactivation |
| 12 | H3pSer10 | Phosphorylation at the 10th Serine residue of the histone H3 | Chromatin | - |
| 13 | H4K16ac | - | Chromatin | A marker of open chromatin |
| 14 | H4K20me1 | Mono-methylation at H4K20 | Chromatin | A marker of transcription inactivation |
| 15 | H4K20me2 | - | Chromatin | A marker of transcription inactivation |
| 16 | H4K20me3 | - | Chromatin | A marker of constitutive-heterochromatin |
| 17 | RNAPIISer5-P | Phosphorylated RNAPII at serine 5 residue | Chromatin | A marker of transcription activation |
| 18 | Fibrillarin | - | Nucleolar | A maker of nucleolar |
| 19 | SF3a66 | Splicing factor 3a subunit 66 | Speckle | A maker of speckle |
| 20 | Lamin B1 | - | Nuclear periphery | A marker of nuclear periphery |

**Table S2.** The criteria of annotating nuclear compartments based on the clusters of gaussian mixture model.

| Annotating steps | Name of Compartment | Abbreviation of interest | Description of Compartment |
| --- | --- | --- | --- |
| 1 | Repeat elements | repeat | Z-score of Minsat > 3 or z-score of Teromere >3 |
| 2 | Nuclear speckle | speckle | Z-score of SF3a66 >3 |
| 3 | Nucleolar | nucleolar | Z-score of Fibrillarin > 1.75 |
| 4 | Constitutive heterochromatin | const-hetero | Z-score of H3K9me3 > 3, z-score of H4K20me3 > 3 or z-score of mH2A1 > 3 |
| 5 | Transcriptionally active compartment-high | active-high | Z-score of RNAPIISer5P > 2. |
| 6 | Transcriptionally mixed compartment | mixed | Z-score of H3K9me3 > 2, z-score of H4K20me3 > 2 or z-score of mH2A1 > 2. |
| 7 | Transcriptionally active compartment-low | active-low | The others. |

**Table S3.** Repeat DNAs that were considered in this study.

| Serial number | Name of repeat DNAs | Abbreviation of repeat DNAs | Description |
| --- | --- | --- | --- |
| 1 | DNA | DNA repeat elements |  |
| 2 | DNA? | DNA repeat elements ? | "?" signifies that "RepeatMasker" was unsure of the classification. |
| 3 | LINE | Long interspersed nuclear elements |  |
| 4 | LINE? | Long interspersed nuclear elements ? | "?" signifies that "RepeatMasker" was unsure of the classification. |
| 5 | LTR | Long terminal repeat elements |  |
| 6 | LTR? | Long terminal repeat elements ? | "?" signifies that "RepeatMasker" was unsure of the classification. |
| 7 | Low_complexity | Low complexity repeats |  |
| 8 | Other | Other repeats |  |
| 9 | RC | Rolling circle |  |
| 10 | RC? | Rolling circle ? |  |
| 11 | RNA | RNA repeats |  |
| 12 | SINE | Short interspersed nuclear elements |  |
| 13 | Satellite | Satellite repeats |  |
| 14 | Simple_repeat | Simple repeats (micro-satellites) |  |
| 15 | Unknown | Unknown |  |
| 16 | rRNA | ribosomal RNA |  |
| 17 | scRNA | small cytoplasmic RNA |  |
| 18 | snRNA | small nuclear RNA |  |
| 19 | srpRNA | signal recognition particle RNA |  |
| 20 | tRNA | transfer RNA |  |

**Table S4.** Estimated periods of each PC mode.

| Chromosome | PCmode1 | PCmode2 | PCmode3 |
| --- | --- | --- | --- |
| chr1 | 660 min | 295 min | 287 min |
| chr2 | 660 min | 303 min | 277 min |
| chr3 | 660 min | 257 min | 239 min |
| chr4 | 660 min | 298 min | 274 min |
| chr5 | 660 min | 295 min | 288 min |
| chr6 | 660 min | 273 min | 266 min |
| chr7 | 660 min | 287 min | 274 min |
| chr8 | 660 min | 289 min | 266 min |
| chr9 | 660 min | 283 min | 239 min |
| chr10 | 660 min | 310 min | 291 min |
| chr11 | 660 min | 299 min | 246 min |
| chr12 | 660 min | 276 min | 257 min |
| chr13 | 660 min | 277 min | 243 min |
| chr14 | 660 min | 295 min | 250 min |
| chr15 | 660 min | 300 min | 286 min |
| chr16 | 660 min | 321 min | 279 min |
| chr17 | 660 min | 353 min | 274 min |
| chr18 | 660 min | 307 min | 259 min |
| chr19 | 660 min | 346 min | 334 min |
| chrX | 660 min | 270 min | 251 min |

